# A hepatocyte-specific cytochrome *c* oxidase deficiency in mice leads to a lymphopenia owing to deficiencies in bone marrow progenitors

**DOI:** 10.1101/2024.08.30.609186

**Authors:** KimAnh T. Pioli, Sampurna Ghosh, Aren Boulet, Scot C. Leary, Peter D. Pioli

## Abstract

Mutations that negatively impact mitochondrial function are highly prevalent in humans and lead to disorders with a wide spectrum of disease phenotypes, including deficiencies in immune cell development and/or function. Previous analyses of mice with a hepatocyte-specific cytochrome *c* oxidase (COX) deficiency revealed an unexpected peripheral blood leukopenia associated with splenic and thymic atrophy. Here, we use mice with a hepatocyte-specific deletion of the COX assembly factor *Sco1* to show that metabolic defects extrinsic to the hematopoietic compartment lead to a pan-lymphopenia represented by severe losses in both B and T cells. We further demonstrate that immune defects in these mice are associated with the loss of bone marrow lymphoid progenitors common to both lineages and early signs of autoantibody-mediated autoimmunity. Our findings collectively point to a significant role for hepatocyte dysfunction as an instigator of immunodeficiency in patients with congenital mitochondrial defects who suffer from chronic or recurrent infections.

## Introduction

Mitochondria are multifaceted organelles that fulfill a diverse number of functions beyond ATP production that are essential to cellular and systemic homeostasis^1–5^. In mammals, organelle function requires the coordinate expression of roughly 1,100 proteins of dual genetic origin^6^. Mutations in these mitochondrially- or nuclear-encoded proteins are a frequent cause of human disease, accounting for roughly 300 distinct disorders with a cumulative estimated birth prevalence of ∼1 in 3,500^7–11^. While mitochondrial disorders range in their presentation from isolated, tissue-specific deficits to multisystem disease^7,8,12^, identifying mechanisms that explain the underlying basis of the observed clinical heterogeneity remains a major challenge.

Most extrinsic and intrinsic inputs that contribute to the clinical heterogeneity of mitochondrial disorders have yet to be identified; however, there is a growing appreciation of immune system involvement in disease etiology in some patient cohorts. Indeed, a recent set of elegant studies firmly established that aberrant leukocyte activity is responsible for many of the neurological and metabolic facets of disease in a mouse model of Leigh syndrome^13,14^. More commonly, however, mitochondrial disease patients like those with Barth or Pearson syndrome present with a lineage-specific or global leukocyte deficiency^15,16^, which makes them susceptible to chronic or recurrent infection^17–19^. It is therefore easy to surmise that immune-related abnormalities in these patients may be driven solely by intrinsic, germline mutations leading to deficiencies in mitochondrial function. Consistent with this idea, hematopoietic stem cells (HSCs) which sit at the top of the immune system hierarchy are particularly sensitive to changes in mitochondrial function^20^. Deletion of *Foxo3* in HSCs, for example, leads to an increased reliance on glycolysis and elevated reactive oxygen species (ROS) production^21^. This altered metabolic profile in turn muted the ability to maintain this HSC population in the bone marrow (BM) and reduced the capacity of these cells to reconstitute the immune system in transplantation experiments^21^. Knockout of another transcription factor, *Srebf1c*, similarly impaired HSC function and transplantation potential by affecting mitochondrial activity and ROS generation^22^. Depletion of the ADP/ATP exchanger *Ant2*, a gene product specific to mitochondria, profoundly impaired erythrocyte and B cell development while largely sparing the myeloid and T cell lineages^23^. Why then immune involvement is absent in most mitochondrial disorders and what dictates whether mutations in nuclear-encoded, mitochondrial gene products with disparate functions affect or spare aspects of immune system function in the context of mitochondrial disease is unclear.

We previously demonstrated that mice with a hepatocyte-specific cytochrome *c* oxidase (COX) deficiency^24^ displayed a significant leukopenia in the blood, and prominent atrophy of the spleen (SPL) and thymus (THY) which play critical roles in B and T cell development, respectively. These phenotypes were met with a profound increase in plasma α-fetoprotein (AFP) levels which contributed to the loss of circulating leukocytes in these mice^24^. However, it was unclear whether other aspects of immune system development and/or function were compromised and contributed to the leukopenia in these mouse models. Here, we further investigate whether functional deficits in the BM, SPL and THY also contribute to the observed immunodeficiency, and find that hepatocyte-specific metabolic dysfunction results in the loss of BM lymphoid progenitors which ultimately gives way to B and T cell developmental abnormalities in the SPL and THY. Rather unexpectedly, these mice present with increased plasma IgA and indicators of altered B cell tolerance leading to antibody-mediated reactivity to liver antigens. Finally, plasma profiling reveals the altered abundance of numerous proteins which are predicted to negatively impact lymphopoiesis.

## Results

### Hepatocyte-specific deletion of Sco1 leads to impaired thymopoiesis and reductions in peripheral T cell populations in the SPL

The THY is the primary site of T cell development, or thymopoiesis, and was shown to prematurely atrophy in mice lacking *Sco1* expression in hepatocytes^24^. To better understand the functional consequences of this observation, we generated hepatocyte-specific *Sco1* knockout mice, hereafter referred to as *Sco1* mice, as previously described^1,2^ and confirmed that they had significantly reduced body weight (**Figure S1A**)^24,25^ and significantly increased plasma AFP levels (**Figure S1B**)^24^ relative to wildtype (*WT*) littermates at postnatal day 47 (P47).

At P47, the *Sco1* THY demonstrated a significant reduction in cellularity regardless of sex (**Figure S1C**). Thymopoiesis requires transit through multiple developmental stages before the release of mature CD4^+^ or CD8^+^ single positive (SP) T cells to the periphery. As such, it is possible that deficiencies at any number of steps could result in THY atrophy. We therefore performed flow cytometry to dissect any alterations in T cell development present in *Sco1* mice. Within the CD4^-^ CD8^-^ double negative (DN) compartment, T cell progenitors lacking surface markers associated with other hematopoietic lineages (Lin^-^) can be identified based upon combinations of CD44 and CD25 (DN1-4) (**Figure 1A**). Within the DN1 population, early THY progenitors (ETPs) that seed the THY express high amounts of CD117 and CD44 (**Figure 1A**). ETPs were significantly decreased in *Sco1* compared to *WT* THY (**Figures 1A-1B**), as were the DN1-4 stages (**Figures 1C-1F**). Impaired development at subsequent stages of T cell maturation including CD4^+^ CD8^+^ double positive (DP) as well as TCRβ^+^ 4SP and TCRβ^+^ 8SP stages was also observed in the P47 *Sco1* THY (**Figures 1G-1J**). Taken together, these data indicate that hepatocyte-specific deletion of *Sco1* leads to an overall reduction in the THY T cell developmental pipeline rather than blockade at a single developmental step. Furthermore, even though female mice display more robust thymopoiesis compared to males at young ages^26,27^, the defects observed in the *Sco1* THY were conserved between sexes.

**Figure 1.**
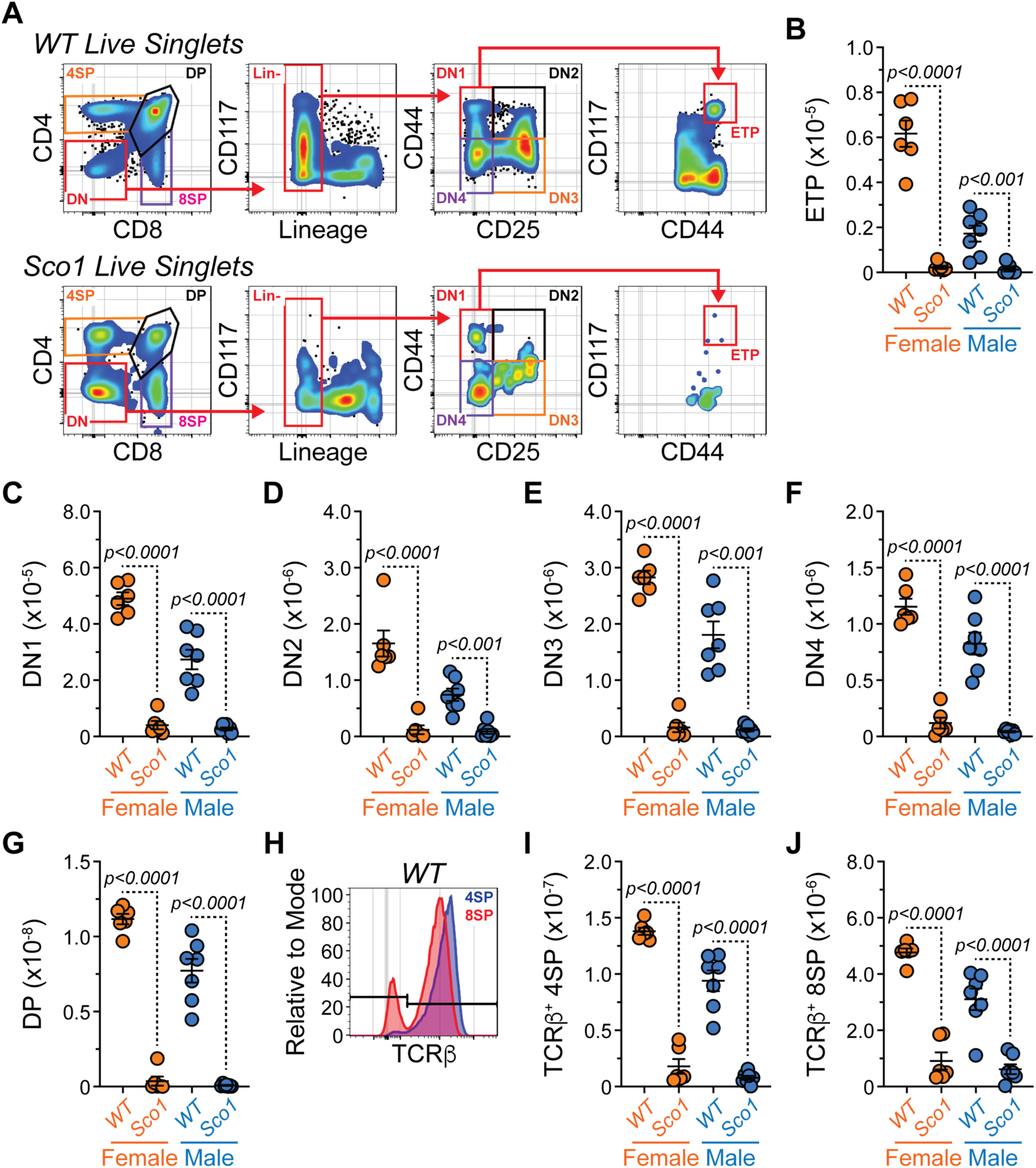
Hepatocyte-specific deletion of *Sco1* leads to impaired thymopoiesis at P47. Related to Figure S1. (A) Flow cytometry plots depicting representative gating of various progenitor and T cell populations within *WT* and *Sco1* THY. Cells pre-gated on total live singlets. (B-G) Numbers of (B) ETPs, (C) DN1, (D) DN2, (E) DN3, (F) DN4 and (G) DP thymocytes from *WT* and *Sco1* THY. (H) Flow cytometry overlays showing TCRβ staining on the surface of 4SP and 8SP cells from *WT* THY. Gates indicate negative and positive staining. (I-J) Numbers of (I) TCRβ^+^ 4SP and (J) TCRβ^+^ 8SP thymocytes from *WT* and *Sco1* THY. (B-G, I-J) Symbols represent individual mice. Horizontal lines represent mean ± SEM. *WT* Female: n = 6, *Sco1* Female: n = 6, *WT* Male: n = 7, *Sco1* Male: n = 7. Statistics: Unpaired Student’s t-Test.

To determine how changes in the THY affected peripheral T cells, we analyzed various T cell populations in the SPL of *WT* and *Sco1* mice. Consistent with previous observations regarding SPL size^24^, the *Sco1* SPL displayed a significant reduction in overall cellularity (**Figure S1D**) that included a decrease in both TCRβ^+^ 4SP and TCRβ^+^ 8SP T cell numbers (**Figures 2A-2C**). Peripheral T cells are a composite population that includes naïve cells yet to encounter antigen and memory T cells such as the central memory (CM) and effector memory (EM) subsets^28^. Using flow cytometry, we could identify all 3 populations within the SPL TCRβ^+^ 4SP compartment (**Figure 2D**) and found that each was significantly reduced in *Sco1* mice (**Figures 2E-2G**). We performed a similar analysis within the SPL TCRβ^+^ 8SP population (**Figure 2H**) and, not surprisingly, observed that naïve (**Figure 2I**), CM (**Figure 2J**) and EM (**Figure 2K**) TCRβ^+^ 8SP cells were also decreased in the P47 *Sco1* SPL. Collectively, these data suggest that diminished T cell numbers are maintained in peripheral lymphoid organs such as the SPL of *Sco1* mice.

**Figure 2.**
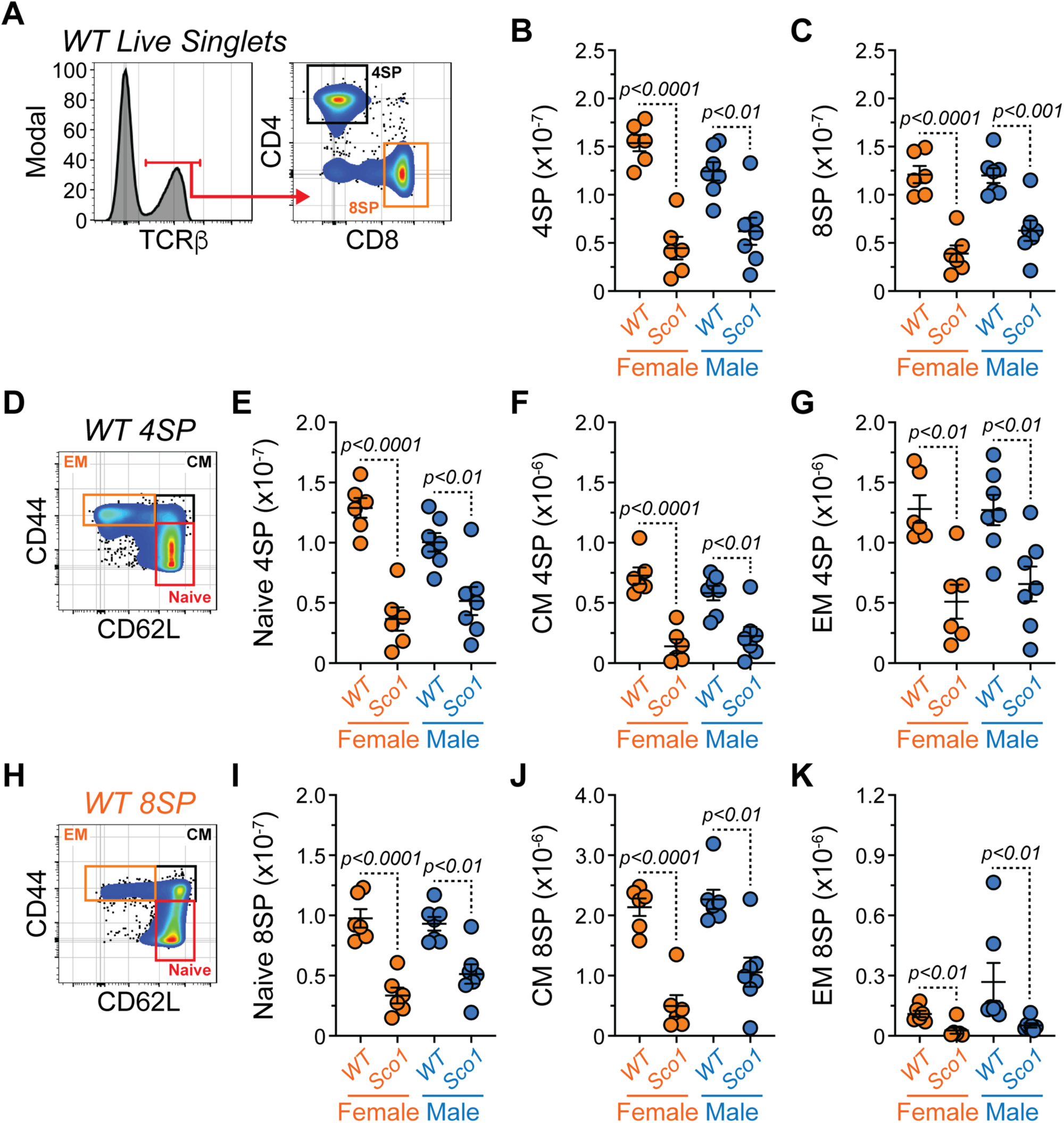
SPL peripheral T cell populations are reduced in P47 *Sco1* mice. Related to Figure S1. (A) Flow cytometry plots depicting representative gating of 4SP and 8SP T cells in *WT* SPL. Cells pre-gated on total live singlets. (B-C) Numbers of (B) 4SP and (C) 8SP T cells from *WT* and *Sco1* SPL. (D) Flow cytometry plots depicting representative gating of naïve, CM and EM 4SP T cells within *WT* SPL. Cells pre-gated on total live singlets. (E-G) Numbers of (E) naïve, (F) CM and (G) EM 4SP T cells from *WT* and *Sco1* SPL. (H) Flow cytometry plots depicting representative gating of naïve, CM and EM 8SP T cells within *WT* SPL. Cells pre-gated on total live singlets. (I-K) Numbers of (I) naïve, (J) CM and (K) EM 4SP T cells from *WT* and *Sco1* SPL. (B-C, E-G, I-K) Symbols represent individual mice. Horizontal lines represent mean ± SEM. *WT* Female: n = 6, *Sco1* Female: n = 6, *WT* Male: n = 7, *Sco1* Male: n = 7. Statistics: Unpaired Student’s t-Test.

### Hematopoietic stem and progenitor cell populations (HSPCs) are altered in Sco1 mice

Defects in thymopoiesis can be caused by intrinsic alterations within the THY, the failure of ETPs to enter the THY or changes in the number or function of upstream progenitors in the BM. To address these disparate possibilities, we performed flow cytometry to evaluate HSPCs in the BM ranging from HSCs to more lineage biased progenitors (**Figure 3A**). In general, *Sco1* BM was hypocellular compared to that of *WT* animals from both sexes (**Figure S1E**). Within the Lin^-^ CD117^+^ Sca-1^+^ (LKS) CD135^-^ compartment (**Figure 3A**), HSCs, multipotent progenitors (MPPs), MPPs with extensive megakaryocyte and erythroid potential (MPP^Mk/E^) as well as MPPs with primarily granulocyte and monocyte potential (MPP^G/M^) were defined as proposed by Challen *et al*^29^. HSCs demonstrated a female-specific reduction in *Sco1* mice (**Figure 3B**). However, MPP, MPP^Mk/E^ and MPP^G/M^ progenitor populations were unaffected in *Sco1* BM of either sex (**Figures 3C-3E**). Similarly, the Lin^-^ CD117^+^ Sca-1^-^ (LKS^-^) population which is collectively comprised of lineage-specified myeloid progenitors was equivalent across genotypes (**Figure 3F**). In contrast, MPPs with significant lymphoid potential (MPP^Ly^) (**Figure 3G**), which can be recognized via CD135 expression within the LKS compartment^29^ (**Figure 3A**) along with downstream common lymphoid progenitors (CLPs)^30^ (**Figures 3H-3I**), were underrepresented in *Sco1* BM of both sexes. As MPP^Ly^ and CLP populations will ultimately contribute to B and T cell production, these data indicate that BM progenitor populations common to B and T cell development are depleted in *Sco1* mice.

**Figure 3.**
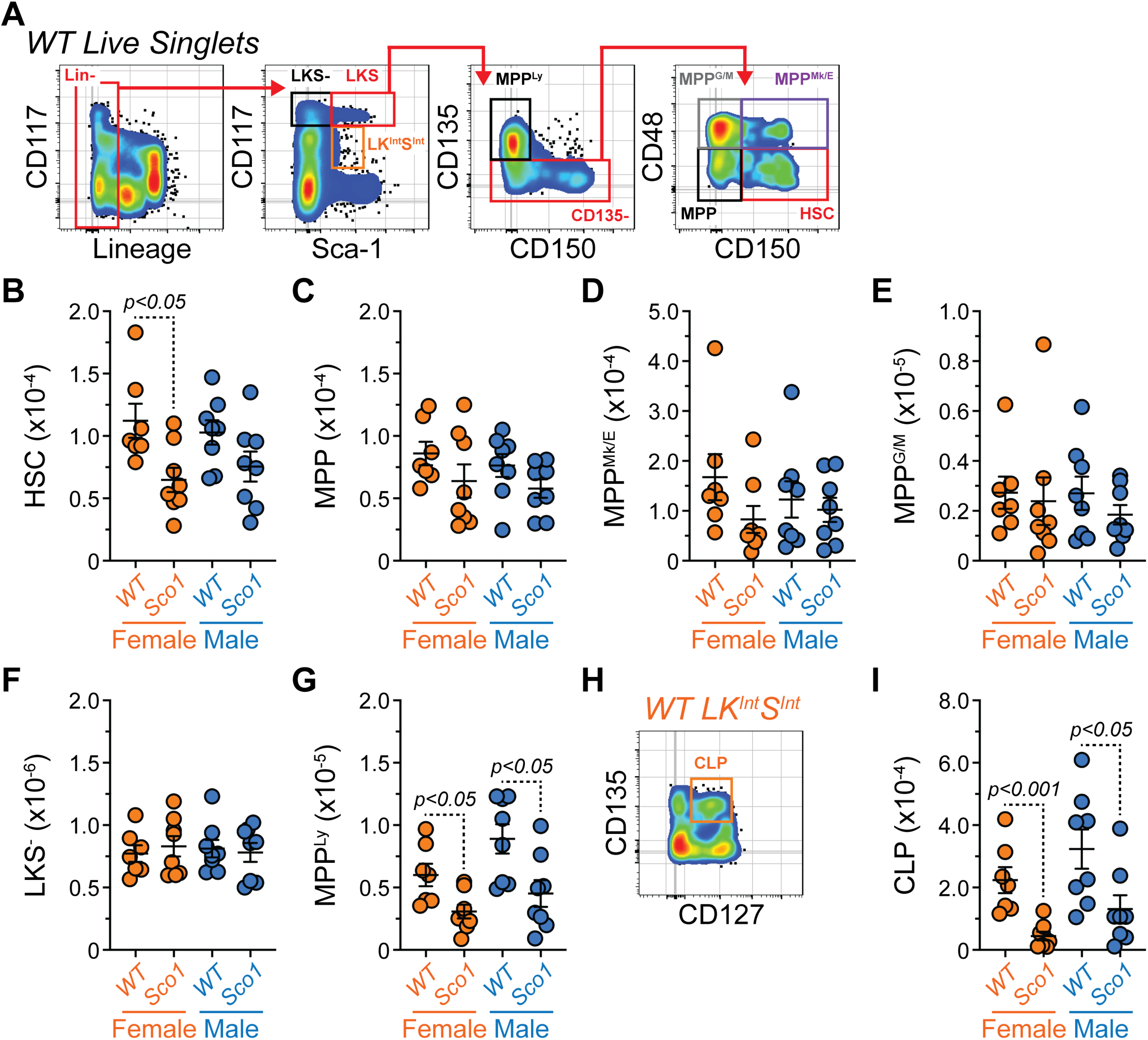
P47 *Sco1* mouse BM displays reduced lymphoid progenitors common to both B and T cell development. Related to Figure S1. (A) Flow cytometry plots depicting representative gating of HSPCs in *WT* BM. Cells pre-gated on total live singlets. (B-G) Numbers of (B) HSC, (C) MPP, (D) MPP^Mk/E^, (E) MPP^G/M^, (F) LKS^-^ and (G) MPP^Ly^ progenitor populations from *WT* and *Sco1* BM. (H) Flow cytometry plot depicting representative gating of CLPs within the LK^Int^S^Int^ population from *WT* BM. (I) Numbers of CLPs from *WT* and *Sco1* BM. (B-G, I) Symbols represent individual mice. Horizontal lines represent mean ± SEM. *WT* Female: n = 7, *Sco1* Female: n = 8, *WT* Male: n = 8, *Sco1* Male: n = 8. Statistics: Unpaired Student’s t-Test.

### Early B cell development is suppressed in the BM of Sco1 mice

Given that we observed alterations in the abundance of select BM progenitors in *Sco1* mice, we next investigated how this impacted downstream lineage output. Within the BM, the 2 major lineages produced are cells of myeloid origin and B cells. Using flow cytometry, we identified the myeloid compartment as CD11b^+^ and further subdivided these cells into monocytes (CD11b^+^ Ly-6C^HI^ Ly-6G^-^) and granulocytes (Ly-6C^-/LO^ Ly-6G^+^) (**Figure 4A**) using previously published methods^30^. BM from both female and male *Sco1* mice displayed a significant decrease in CD11b^+^ myeloid cells (**Figure 4B**), while only male *Sco1* BM exhibited a clear reduction in both monocyte (**Figure 4C**) and granulocyte (**Figure 4D**) populations.

**Figure 4.**
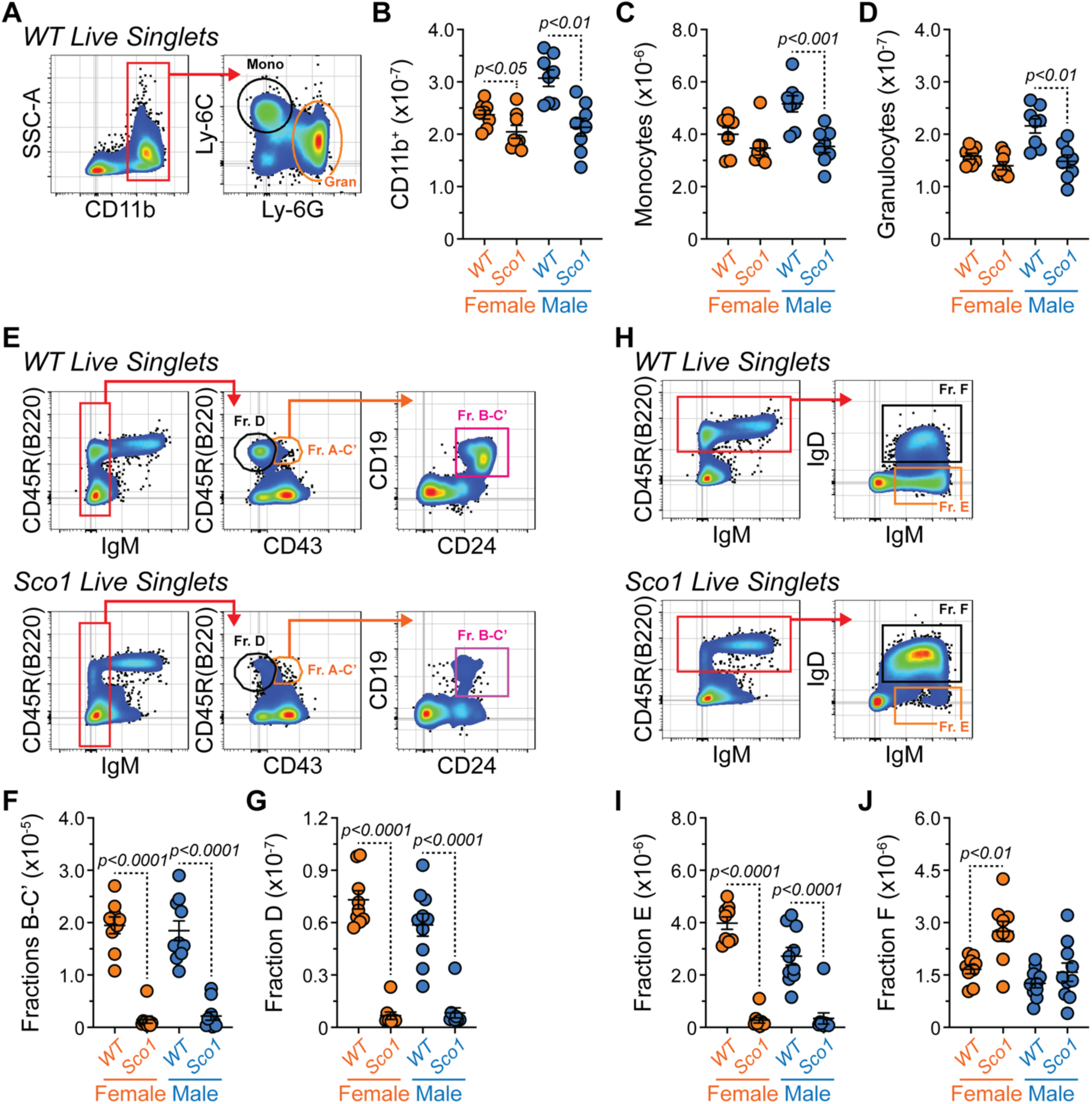
BM B cell development is severely depleted in P47 *Sco1* mice. Related to Figures S1-S2. (A) Flow cytometry plots depicting representative gating of myeloid populations in *WT* BM. Cells pre-gated on total live singlets. (B-D) Numbers of (B) total CD11b^+^ myeloid cells, (C) monocytes and (D) granulocytes from *WT* and *Sco1* BM. (E) Flow cytometry plots depicting representative gating of B cell Hardy Fractions A-D within *WT* and *Sco1* BM. Cells pre-gated on total live singlets. (F-G) Numbers of (F) Hardy Fractions B-C’ and (G) Hardy Fraction D B cells from *WT* and *Sco1* BM. (H) Flow cytometry plots depicting representative gating of B cell Hardy Fractions E-F within *WT* and *Sco1* BM. (I-J) Numbers of (I) Hardy Fraction E and (J) Hardy Fraction F B cells from *WT* and *Sco1* BM. (B-D, F-G, I-J) Symbols represent individual mice. Horizontal lines represent mean ± SEM. Statistics: Unpaired Student’s t-Test. (B-D) *WT* Female: n = 8, *Sco1* Female: n = 8, *WT* Male: n = 8, *Sco1* Male: n = 8. (F-G, I-J) *WT* Female: n = 9, *Sco1* Female: n = 9, *WT* Male: n = 10, *Sco1* Male: n = 10.

To evaluate B cell development, we employed a modified version of the Hardy fraction scheme^31^. Within the IgM^-^ BM, Fractions A-C’ were identified as CD45R(B220)^+^ CD43^+^ (**Figure 4E**). Subsequently, committed B cell progenitors (Fractions B-C’) were gated as CD19^+^ CD24^+^ (**Figure 4E**). Overall, numbers of Fractions B-C’ were reduced in *Sco1* BM (**Figure 4F**). This compartment includes early Pro-B cells (Fraction B) as well as late Pro-B and large Pre-B cells (Fractions C-C’) which are actively cycling^31^. Using forward scatter (FSC-A) (i.e., cell size) as a surrogate for proliferation (**Figure S2A**), we observed a decrease in this parameter within cells from *Sco1* Fractions B-C’ (**Figures S2A-S2B**). Based upon Ly-51(BP-1) expression^31^ (**Figure S2C**), we further delineated between Fractions B and C-C’ and observed that both were decreased in *Sco1* BM (**Figures S2D-S2E**). Following VDJ recombination of the immunoglobulin heavy chain, validation of the Pre-B cell receptor (Pre-BCR) and exit from the cell cycle, B cell progenitors transition to the small Pre-B stage (Fraction D) (**Figure 4E**) whereupon they undergo recombination of the immunoglobulin light chain^31^. Similar to upstream Fractions, Fraction D was also reduced in *Sco1* mice (**Figure 4G**). If VDJ recombination was non-functional, then *Sco1* mice would lack B cells expressing surface immunoglobulins/BCRs. To address this possibility, we evaluated IgM and IgD surface expression on CD45R(B220)^+^ cells in the BM (**Figure 4H**). IgM^+^ Fraction E cells represent the immediate descendants from the Small Pre-B Fraction D and were underrepresented in *Sco1* BM (**Figure 4I**). However, cells expressing both IgM and IgD (Fraction F) were present at normal or increased amounts in *Sco1* BM (**Figure 4J**). Since Fraction F represents mature B cells that recirculate to the BM, these data indicate that while the B cell developmental pipeline as a whole is significantly reduced in *Sco1* BM, it is not blocked at any single stage of maturation.

To further confirm the reduced presence of B cell progenitors in *Sco1* BM, we performed *in vitro* Pre-B colony assays. For these experiments, 10^5^ total cells from P47 *WT* and *Sco1* BM were plated in 12-well dishes in the presence of 10 ng/mL interleukin (IL)-7 (**Figure S2F**). After 10 days, macroscopic colonies were counted in a blinded manner and cellular identity was confirmed by flow cytometry. These analyses revealed lower colony formation with *Sco1* than *WT* BM (**Figure S2G**) which, based on the reduction in CD19 geometric mean fluorescence intensity (gMFI) (**Figures S2H-S2I**), may be attributable to a diminished contribution of the B cell lineage. Notably, impaired colony formation appeared to impact the female sex to a greater degree.

### Immature and mature B cell populations are reduced in the Sco1 SPL

After IgM-expressing B cells exit the BM, they immigrate to the SPL and undergo additional maturation steps which include the immature transitional B cell (T1 and T2/T3) stages as well as the mature follicular (FO) and marginal zone (MZ) stages (**Figure 5A**). For the flow cytometric analysis of these populations, CD138^HI^ cells were excluded to prevent any numerical bias generated by contaminating antibody-secreting cells (ASCs) (see below section) (**Figure 5A**). As might be expected based upon the BM data, the *Sco1* SPL possessed a significant decrease in the T1 (**Figure 5B**), T2/T3 (**Figure 5C**), FO (**Figure 5D**) and MZ (**Figure 5E**) B cell populations.

**Figure 5.**
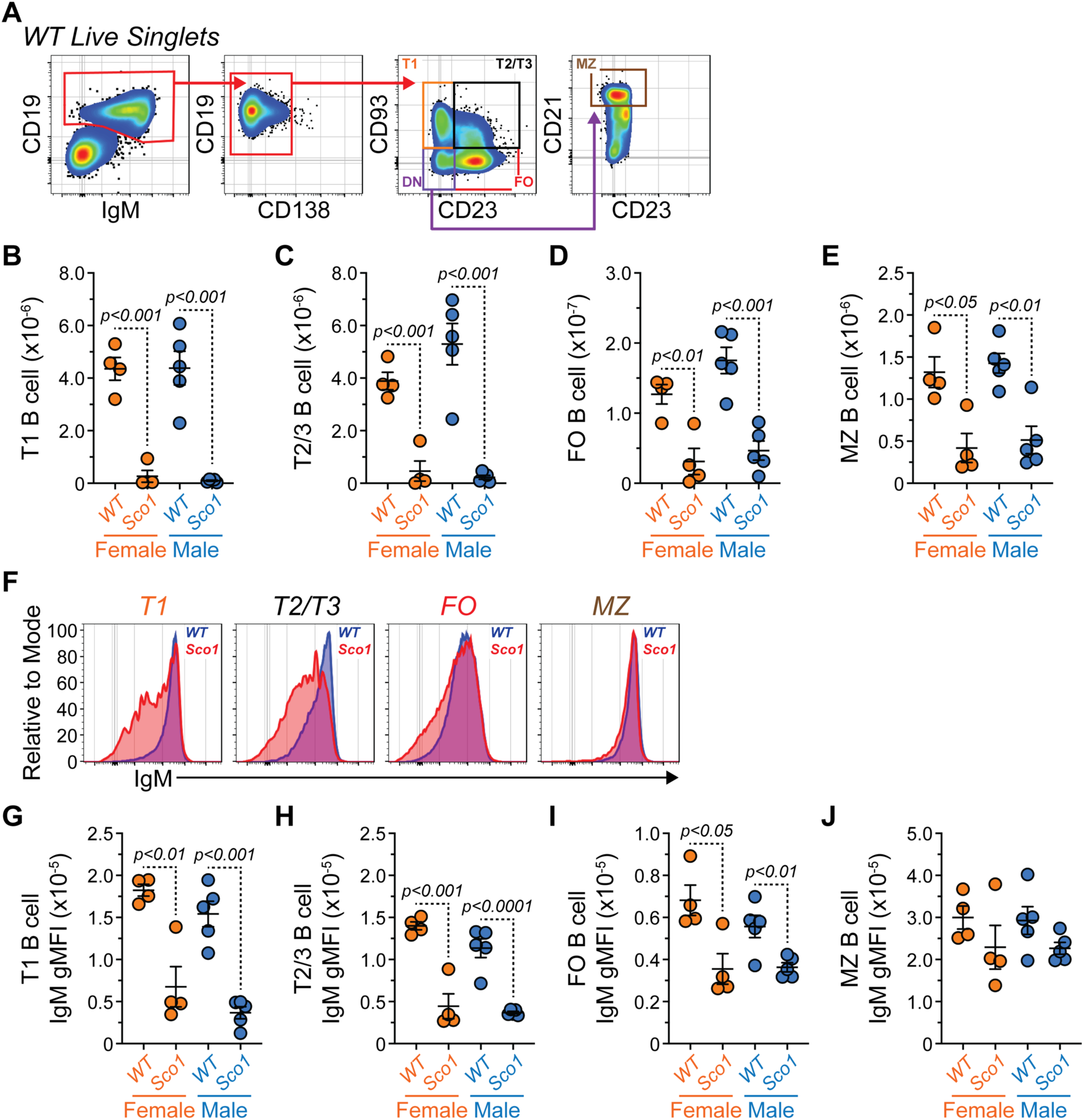
SPL B cell maturation is dysfunctional in P47 *Sco1* mice. Related to Figures S1 and S3. (A) Flow cytometry plots depicting representative gating of B cell populations in *WT* SPL. Cells pre-gated on total live singlets. DN = double negative. (B-E) Numbers of (B) T1, (C) T2/3, (D) FO and (E) MZ B cells from *WT* and *Sco1* SPL. (F) Flow cytometry overlays showing IgM staining on the surface of T1, T2/3, FO and MZ B cells from *WT* and *Sco1* SPL. (G-J) IgM surface staining gMFI for (G) T1, (H) T2/3, (I) FO and (J) MZ B cells from *WT* and *Sco1* SPL. (B-E, G-J) Symbols represent individual mice. Horizontal lines represent mean ± SEM. *WT* Female: n = 4, *Sco1* Female: n = 4, *WT* Male: n = 5, *Sco1* Male: n = 5. Statistics: Unpaired Student’s t-Test.

Similar to developing T cells, B cells also undergo checkpoints to ensure their lack of self-reactivity. In the SPL, this takes place at the transitional stages and can lead to downregulation of IgM in an attempt to nullify the responsiveness (i.e., make anergic) of autoreactive B cells^32^. Along these lines, we observed downregulation of surface IgM (**Figure 5F**) which was restricted to T1 (**Figure 5G**), T2/T3 (**Figure 5H**) and FO (**Figure 5I**) *Sco1* B cells while MZ B cells in *Sco1* mice were relatively unaffected (**Figure 5J**). IgM and IgD are derived from the same RNA due to alternative splicing and the function of Zfp318^33,34^. While they both recognize the same antigen, IgM and IgD are functionally distinct with IgD being more tolerogenic *in vivo*^35^. Examination of IgD surface expression across the various B cell subsets (**Figures S3A-S3E**) revealed a consistent increase at the T2/3 stage in both female and male *Sco1* B cells (**Figure S3C**). Additionally, there were some notable sex differences as T1 and MZ *Sco1* B cells appeared to upregulate IgD in a female-specific manner (**Figures S3B and S3E**). More recent work has shown intermediate expression of CD138 to correlate with the induction of B cell anergy^36^ and the levels of this protein can be modulated by BCR signaling^36,37^. Accordingly, shifts in CD138 surface expression were observed in SPL *Sco1* B cells (**Figures S3F-S3J**) as represented by increased CD138 in T1 (**Figure S3F**) and MZ (**Figure S3J**) B cells and decreased CD138 in FO B cells (**Figure S3I**) from *Sco1* mice. These data suggest that B cells developing in mice with hepatocyte-specific deletion of *Sco1* undergo shifts in their BCRs which may influence their reactivity to self-antigen.

### Sco1 mice have reduced BM antibody-secreting cells (ASCs) but elevated plasma IgA levels

To better understand how alterations in B and T cell development influenced immunity in *Sco1* mice, we evaluated populations generated downstream of B cell activation. In this instance, we focused on ASCs which are terminally differentiated B cells whose canonical role is to produce protective antibodies^38^. Using flow cytometry, ASCs were identified as CD138^HI^ CD90.2^-^ CD44^+^ (**Figure 6A**). Examination of these cells in the P47 SPL showed a trend towards reduced numbers in *Sco1* females with little impact in *Sco1* males (**Figure 6B**). Following their production in the periphery, ASCs migrate to the BM where they can adopt a long-lived phenotype. In both sexes, the P47 *Sco1* BM showed a significant decrease in ASCs (**Figure 6C**). ASCs can be generated via multiple routes including spontaneous germinal center reactions that take place in the absence of overt infection or immunization^39^. As such, we wanted to determine if SPL germinal center B cells (GCBs) were reduced in *Sco1* mice and if this correlated with the observed changes in ASC numbers. GCBs (CD19^+^ CD138^-/INT^ CD95(Fas)^+^ GL7^+^) were present in the SPLs of both genotypes (**Figure S4A**). As a percentage of total B cells, GCBs were equivalent between *WT* and *Sco1* mice suggesting normal regulation of this population (**Figure S4B**). However, absolute numbers of these cells were significantly decreased in female *Sco1* animals (**Figure S4C**). While the male sex also displayed reduced *Sco1* SPL GCBs, mouse-to-mouse variability likely prevented this trend from reaching statistical significance (**Figure S4C**).

**Figure 6.**
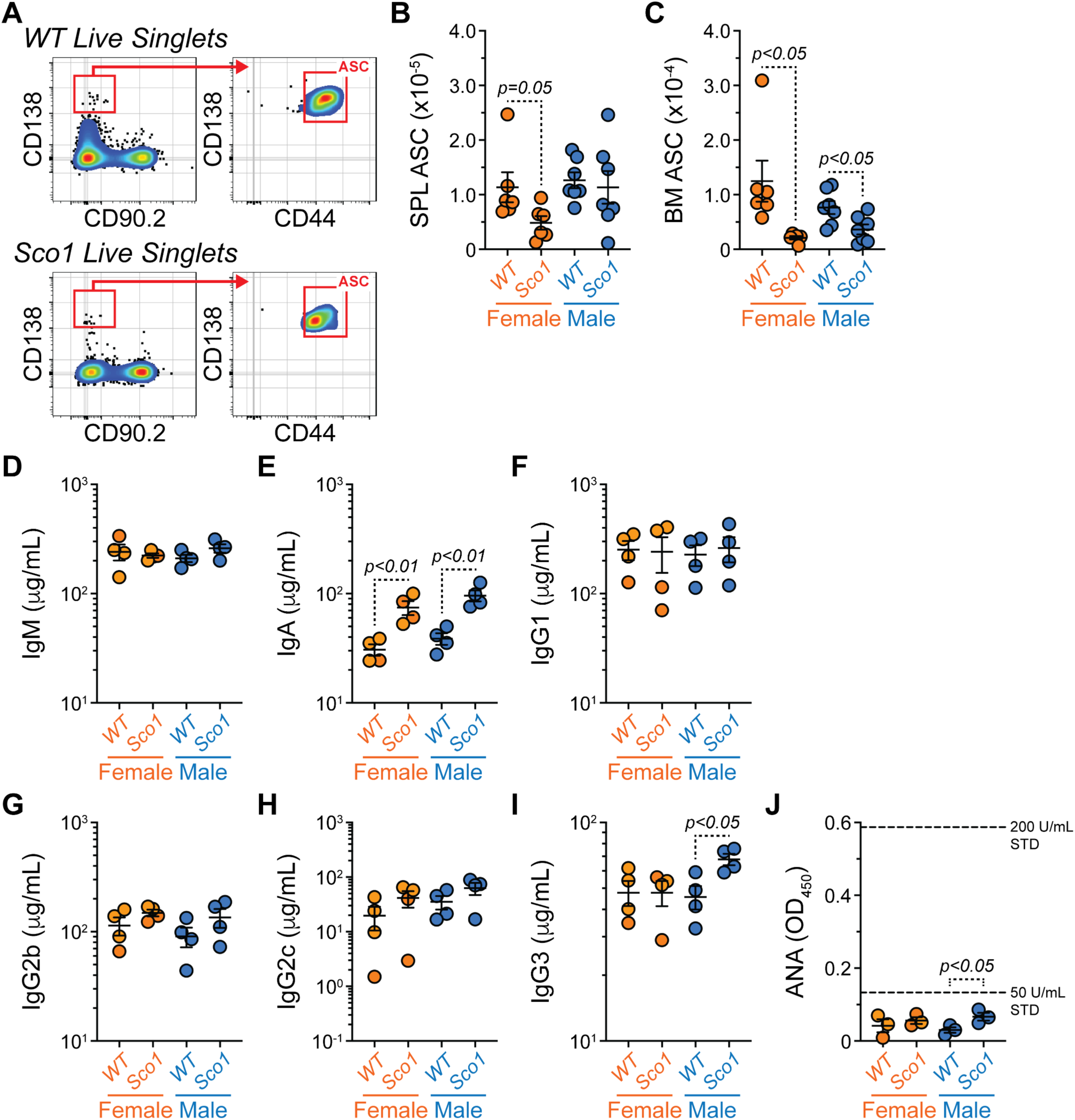
ASC populations are altered in P47 *Sco1* mice which is accompanied by increased plasma IgA. Related to Figures S1 and S4. (A) Flow cytometry plots depicting representative gating of ASCs in *WT* and *Sco1* SPL. Cells pre-gated on total live singlets. (B-C) Numbers of (B) SPL ASCs and (C) BM ASCs from *WT* and *Sco1* mice. (D-I) Levels of (D) IgM, (E) IgA, (F) IgG1, (G) IgG2b, (H) IgG2c and (I) IgG3 in plasma from *WT* and *Sco1* mice. (J) ANA levels in *WT* and *Sco1* plasma as measured by absorbance at OD_450_ using a 1:100 dilution. (B-J) Symbols represent individual mice. Horizontal lines represent mean ± SEM. Statistics: Unpaired Student’s t-Test. (B-C) *WT* Female: n = 6, *Sco1* Female: n = 6, *WT* Male: n = 7, *Sco1* Male: n = 7. (D-I) *WT* Female: n = 4, *Sco1* Female: n = 4, *WT* Male: n = 4, *Sco1* Male: n = 4. (J) *WT* Female: n = 3, *Sco1* Female: n = 3, *WT* Male: n = 3, *Sco1* Male: n = 3.

Next, we correlated SPL GCB and ASC numbers from each animal. The correlation (R^2^) of SPL GCBs and ASCs increased from 0.55 (p<0.05) in *WT* mice (**Figure S4D**) to 0.83 (p<0.0001) in *Sco1* animals (**Figure S4E**), indicating a stronger relationship between the SPL germinal center reaction and SPL ASC output in the *Sco1* background. Examination of *WT* BM ASCs showed a correlation of 0.62 (p<0.01) with *WT* SPL ASCs (**Figure S4F**). In sharp contrast, the correlation between *Sco1* BM ASCs and *Sco1* SPL ASCs was not significant (R^2^=0.23, p<0.10) revealing a poor relationship between these populations and suggesting that BM ASCs in *Sco1* mice were mostly derived from other organ sites (e.g., intestine).

Based on the above observations, we suspected that plasma antibody levels may be deficient in *Sco1* mice. However, enzyme-linked immunosorbent assays (ELISAs) revealed that IgM was equivalent across genotypes (**Figure 6D**) while plasma IgA was significantly increased in both sexes of P47 *Sco1* mice (**Figure 6E**). Examination of IgG subtypes showed that IgG1 (**Figure 6F**), IgG2b (**Figure 6G**) and IgG2c (**Figure 6H**) were unchanged, while IgG3 exhibited a significant increase specific to *Sco1* male mice (**Figure 6I**). These results collectively indicate that even with reduced B cell numbers, P47 *Sco1* mice do not suffer from an antibody deficiency. Rather, P47 *Sco1* mice possess increased plasma IgA which is consistent with previous reports from human patients with various forms of liver disease^40,41^. Given the increased plasma IgA levels (**Figure 6E**) and the downregulation of surface IgM expression by SPL B cells (**Figures 5G-5I**), we hypothesized that P47 *Sco1* mice possessed elevated levels of plasma autoantibodies (autoAbs) which could target universal (e.g., DNA) or tissue-specific (e.g., soluble liver antigen) autoantigens^42^. To test for the presence of autoAbs against universal autoantigens, we employed a commercial mouse anti-nuclear autoantibody (ANA) testing kit. While there was a slight elevation in male *Sco1* relative to *WT* plasma, ANA abundance in all samples from both sexes was below the quantification limit suggesting an overall lack of this autoAb species in *Sco1* plasma (**Figure 6J**). To determine the presence of autoAbs reactive to the liver, we adapted a previously published flow cytometry method^43^ and tested the ability of *WT* and *Sco1* plasma to bind total liver cell suspensions that had been fixed and permeabilized (**Figures 7A-7B**). As a result of permeabilization, autoAbs in the plasma could bind to cell surface antigens as well as those restricted to the intracellular compartment. After subtracting Igκ+λ-FITC (2° only) background, plasma from *WT* and *Sco1* female mice did not show a clear difference in reactivity to liver cells (**Figures 7C-7D**). This result was consistent whether the data were assessed as % of binding (**Figure 7C**) or as gMFIs indicative of overall staining intensity (**Figure 7D**). In contrast, male *Sco1* plasma possessed increased reactivity to liver antigens compared to male *WT* plasma (**Figures 7E-7F**).

**Figure 7.**
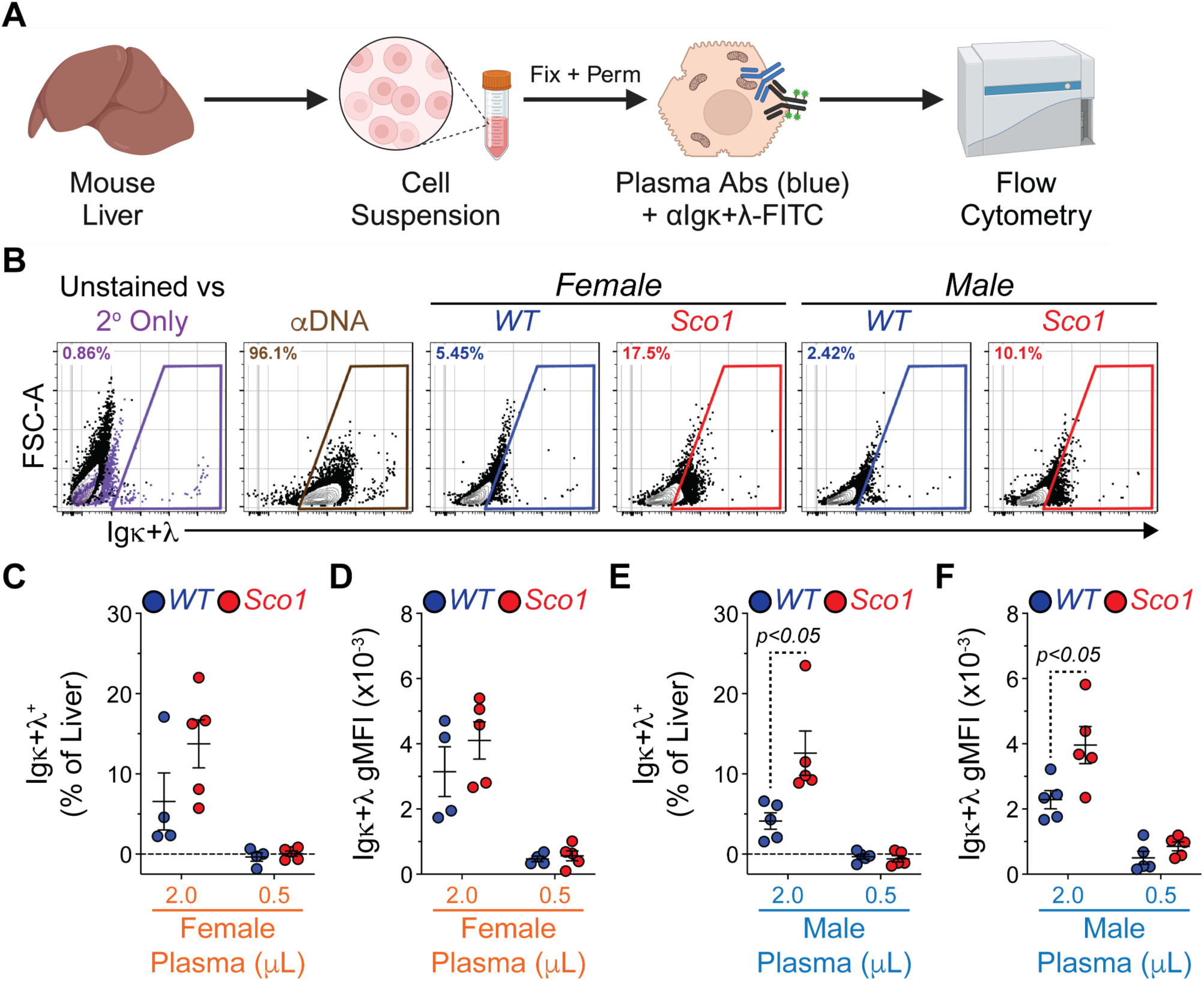
Plasma from P47 *Sco1* mice is autoreactive to mouse liver cell suspensions. (A) Schematic depicting the assay to detect *WT* and *Sco1* plasma autoreactivity to cells isolated from *WT* mouse liver. Figure made with BioRender. (B) Representative flow cytometry plots depicting FSC-A versus Igκ+α staining for liver cells that were unstained, stained with Igκ+α-FITC (2° only) and stained with αDNA positive control antibody or various female and male *WT* and *Sco1* plasma samples in conjunction with Igκ+α-FITC. Boxed areas indicate positive staining over the predominant Igκ+α-FITC (2° only) background signal. %s in plots indicate proportion of liver cells positively stained in each sample. Cells pre-gated on total live singlets. (C-D) Igκ+α-FITC reactivity of female *WT* and *Sco1* plasma presented as (C) % of liver cell binding and (D) gMFI. (E-F) Igκ+α-FITC reactivity of male *WT* and *Sco1* plasma presented as (E) % of liver cell binding and (F) gMFI. (C-F) Values plotted represent those following Igκ+α-FITC (2° only) background subtraction. Symbols represent individual mice. Horizontal lines represent mean ± SEM. *WT* Female: n = 4, *Sco1* Female: n = 5, *WT* Male: n = 5, *Sco1* Male: n = 5. Statistics: Unpaired Welch’s t-Test.

### P47 Sco1 plasma has altered abundance of multiple cytokines with hematopoietic regulatory potential

Elevated AFP levels have previously been associated with the ability of *Sco1* plasma to promote leukocyte apoptosis *in vitro*^24^. As such, we wanted to determine if the abundance of other proteins with putative hematopoietic modulatory function was also altered in P47 *Sco1* plasma. We therefore isolated plasma from *WT* and *Sco1* littermates and generated pools of 4 animals (2 females, 2 males) per genotype. Protein abundance was subsequently assessed using the RayBiotech Mouse L308 antibody array, and considered to be significantly different between the 2 plasma genotypes based on 3 criteria: 1) the protein abundance was regulated in the same direction for all 3 experiments (i.e., Up in *Sco1* or Down in *Sco1*), 2) the average non-transformed fold change (*Sco1*/*WT*) was >2.0 or <0.5 and 3) the standard deviation was <3 for the fold change from all 3 experiments. The resulting analysis identified 23 proteins whose abundance increased in *Sco1* plasma and 133 proteins whose levels decreased in *Sco1* plasma relative to *WT* samples (**Table S1**). As a validation of our analysis, *Sco1* plasma had elevated levels of serum amyloid A1 (SAA1) which is increased in humans with various forms of liver disease^44–46^ as well as mouse models of non-alcoholic fatty liver disease (NAFLD)^45,47^. Additional cytokines related to liver disease or pathology such as thymus and activation-regulated cytokine (TARC)^48^, osteopontin (OPN)^49^, IL-22^50^, IL-17D^51^, IL-17C^52,53^, tumor necrosis factor-alpha (TNF-α)^54^ and receptor activator of nuclear factor κB ligand (RANKL)^55^ (**Figure S5A and Table S1**) were also enriched in *Sco1* plasma. Notably, many of these factors regulate aspects of hematopoiesis. For example, TNF-α influences the behavior of hematopoietic progenitors^30,56,57^ and is a potent repressor of B cell development^58,59^. Furthermore, RANKL^60^ and OPN^61^ regulate the BM HSC niche and OPN acts to limit the overall size of the HSC pool^61^.

In contrast, lymphopoiesis promoting factors such insulin-like growth factor 1 (IGF-1)^62–64^, IL-7^65–67^ and stromal cell-derived factor 1 (SDF-1)/CXCL12^68^ were decreased in *Sco1* plasma (**Figure S5B and Table S1**). Reductions in granulocyte-macrophage colony stimulating factor (GM-CSF)^69^ and macrophage colony stimulating factor (M-CSF)^70^ (**Figure S5B and Table S1**) were similarly observed and may be relevant to the mildly reduced myelopoiesis in the BM of *Sco1* mice (**Figures 4B-4D**). Erythropoietin (EPO) and thrombopoietin (TPO) were also less abundant in *Sco1* plasma (**Figure S5B and Table S1**). Although unrelated to lymphoid and myeloid development, EPO promotes erythropoiesis^71^ while TPO is largely liver-derived and is essential to normal platelet formation^72^ and maintenance of the HSC compartment^73,74^. Further related to HSCs, transforming growth factor-beta 2 (TGF-β2) was the most reduced protein in *Sco1* plasma (**Figure S5B and Table S1**) which is interesting given its importance in regulating early HSPC function^75^. Unrelated to BM hematopoiesis, T helper 2 (T_H_2)-associated cytokines such as IL-13, IL-5 and IL-4 as well as the pan-immunosuppressive cytokine IL-10 were also deficient in *Sco1* plasma (**Figure S5C and Table S1**), indicative of an altered immune response.

To put the altered *Sco1* plasma protein signature into a more global context, we converted protein names to their corresponding gene identities and performed gene ontology (GO) analysis using Metascape^76^ (**Table S2**). Proteins with increased abundance in *Sco1* plasma were significantly associated with GOs related to c-Jun N-terminal kinase (JNK) signaling, the inflammatory response and the migration of leukocytes and epithelial cells (**Figure S5D and Table S2**). GOs associated with a relative decrease in *Sco1* plasma proteins again were correlated with aspects of cell migration as well as pathways regulating cytokine signaling and the immune response (**Figure S5B and Table S2**). Overall, the data indicate that hepatocyte-specific deletion of *Sco1* results in the formation of a circulating cytokine milieu that is predicted to suppress normal immune system function and immune cell development, especially that of B and T cells.

## Discussion

In this study, we show that mice with a hepatocyte-specific COX deficiency develop substantial defects in B and T cell development which correlate with a significant reduction in MPP^Ly^s and CLPs, BM progenitor stages common to both lineages. In contrast, development of myeloid cells in the BM is only mildly affected. We further show that *Sco1* mice possess indicators suggestive of a breach in immune tolerance which include decreased surface IgM on B cells, increased plasma IgA and enhanced reactivity to liver antigens. Finally, we show that these immune phenotypes correlate with the increased abundance of numerous pro-inflammatory factors in the plasma of *Sco1* animals.

*Sco1* is a nuclear-encoded, mitochondrial protein with essential roles in copper delivery to COX during its assembly and the regulation of cellular copper homeostasis^77–80^. In humans, mutations in *SCO1* are exceedingly rare and are associated with highly variable clinical phenotypes tied to neonatal or pediatric mortality^80–82^. A compound heterozygous *SCO1* patient with a nonsense mutation on the paternal allele and a P174L amino acid substitution on the maternal allele exhibited a severe, isolated COX deficiency and died from liver failure and ketoacidotic coma^80^. A patient from a second *SCO1* pedigree harbored a homozygous G132S substitution that resulted in an isolated COX deficiency associated with a multisystem disorder affecting heart, liver and brain function^82^. A third *SCO1* patient possessing a M294V substitution and a frameshift mutation resulting in a premature stop codon (Val93*) succumbed from brain failure in the absence of any apparent liver or heart involvement^81^. Notably, affected tissues from all *SCO1* pedigrees characterized to date also exhibited a severe copper deficiency^78,81,82^. The present study in which loss of *Sco1* expression in hepatocytes leads to a profound COX and copper deficiency in the liver^25^ therefore mirrors the general human condition but most closely resembles the collective molecular phenotypes of the P174L patient.

Our analysis of the *Sco1* BM shows a modest reduction in HSCs specific to females (**Figure 3B**) and a considerable loss of MPP^Ly^s and CLPs common to both sexes (**Figures 3G**, **3I**). As MPP^Ly^s and CLPs are shared progenitors in B and T cell development^83^, it is not surprising that subsequent B cell developmental stages in the BM and SPL as well as T cell development in the THY are greatly reduced. Notably, the abundance of multiple inflammatory cytokines is increased in the plasma of *Sco1* mice (**Table S1 and Figure S5A**), and many of these factors are able to directly suppress B cell development (e.g., TNF-α^58,59^). In contrast, factors known to promote lymphoid development such as IGF-1^62–64^, IL-7^65–67^ and CXCL12^62–64^ are reduced in *Sco1* plasma (**Table S1 and Figure S5B**). While a comparable analysis of these cytokines remains to be done in other *WT* and *Sco1* compartments like the BM or THY, it is quite possible that multiple independent molecular mechanisms coalesce to suppress lymphoid development. Consistent with this idea, we corroborated (**Figure S1B**) that *Sco1* mouse plasma has increased levels of AFP^24^, which we previously showed is capable of promoting apoptosis of peripheral blood leukocytes^24^ and points to a potential mechanism for immunodeficiency in patients with mitochondrial diseases. However, an obvious question remains: does AFP play an active role in the BM deficiencies that we observe herein? AFP has been shown to bind CCR5 on the surface of human monocyte-derived macrophages^84^, and its ability to bind copper is key to AFP-mediated apoptosis of peripheral leukocytes *in vitro*^24^. Studies using zebrafish demonstrated that copper overload impairs proper HSPC proliferation^85^, a phenotype that was also observed in the K562 cell line derived from a patient with chronic myeloid leukemia^85^. While it is unclear if the same effect would translate to committed B and T cell progenitors, many of these populations are highly proliferative and we observe a reduction in cell size, a surrogate for proliferation, in the *Sco1* B cell progenitor Hardy Fractions B-C’ (**Figures S1A-S1B**). Although we have not directly measured CCR5 expression on the surface of these cells, *Ccr5* expression can be induced in mouse HSPCs and immune cells following exposure to total body irradiation^86^ or cytokines such as IL-12^87^, respectively. Of course, the ability of AFP to regulate BM hematopoiesis would also be contingent on its presence in that compartment, either via the bloodstream or a localized increase in expression as seen in BM samples from patients with hepatocellular carcinoma^88,89^. Overall, the data indicate that the lymphopenia observed in *Sco1* mice is due to a multi-hit model in which hematopoietic precursors as well as mature immune cells are targeted.

As might be expected based upon the *Sco1* BM and THY phenotypes (**Figures 3G**, **3I, 4F-4G and 4I-4J**), *Sco1* mice possess a severe SPL lymphopenia consisting of deficiencies in both T and B cells (**Figures 2 and 5**). Naïve, CM and EM subsets are present in CD4 and CD8 T cell populations from the SPL of *Sco1* mice; however, all populations display reductions in their overall cell numbers suggesting an absence of homeostatic proliferation^90,91^. As IL-7 is a critical regulator of this process^91^, the low relative abundance of this cytokine in *Sco1* plasma (**Table S1 and Figure S5B**) may partially explain the observed SPL T cell phenotypes. *Sco1* mice display significant reductions in SPL T1, T2/3, FO and MZ B cell subsets (**Figures 5B-5E**). Unexpectedly, *Sco1* T1, T2/3 and FO B cells possess decreased amounts of surface IgM (**Figures 5G-5I**) which is commonly associated with induction of B cell anergy to suppress the activation of B cells possessing highly autoreactive BCRs^32^. Along these lines, T1 B cells from *Sco1* mice have heightened levels of CD138 which can be modulated by BCR stimulation^36,37^. The T1 B cell stage represents a key peripheral tolerance checkpoint^92^, and these cells are inappropriately expanded in lupus patients^93^. Notably, interferon signaling can enhance the sensitivity of this population to Toll-like receptor (TLR) 7 stimulation^94^ and T1 B cells from TLR7 transgenic mice have an enhanced potential to differentiate into ASCs *in vitro*^95^. Along these lines, *Sco1* mice do not show a loss of plasma antibodies (**Figures 6D**, **6F-6H**) and in fact harbor increased IgA in both sexes (**Figure 6E**) as well as elevated IgG3 in male mice (**Figure 6I**). These observations led us to test for the presence of autoAbs in the plasma of *Sco1* mice. Unlike hepatic autoimmune diseases like hepatitis type 1 and primary biliary cholangitis^42,96^, ANA levels are very low in *Sco1* plasma and mirror those seen in the *WT* background (**Figure 6J**). In contrast, *Sco1* plasma exhibits reactivity to liver antigens (**Figures 7C-7F**). While this observation is largely restricted to males, it is conceivable that the observed sex bias simply reflects a higher degree of variability in this response in females (**Figures 7C-7D**). Alternatively, this variability may reflect the fact that we analyzed mice at P47 owing to the early mortality tied to their liver dysfunction^25^, and that autoAb generation would be detected in older animals of both sexes. While we have not identified the target autoantigen(s), these data raise the possibility that exposure to liver-specific proteins or common non-nuclear molecules derived from hepatocytes may serve as inducing agents of autoAb production in *Sco1* mice. Consistent with this idea, recent work showed that attenuated COX14 function in hepatocytes led to the release of mitochondrial mRNAs into the cytoplasm which then promoted an interferon response^97^. These mitochondrial transcripts, or even mitochondrial DNA or proteins, may also gain access to the cell periphery via necrosis^98^, where they would stimulate circulating mitochondria-specific B cells to differentiate into autoAb-producing ASCs^99^.

Mice with hepatocyte-specific deletion of *Sco1* were originally generated to assess how mitochondrial defects in hepatocytes could contribute to larger, multisystem diseases observed in humans with *SCO1* mutations. However, it is interesting to note that the elevated IgA and presence of autoAbs observed here are reminiscent of phenotypes seen in alcoholic liver disease^100^ and various forms of autoimmune hepatitis^101^. Furthermore, mitochondrial dysfunction is associated with a number of acquired chronic liver disorders^102^. The continued investigation of *Sco1* mice therefore will allow for a greater understanding of how liver defects in mitochondrial function lead to immune system alterations, without the caveats related to disease modeling that accompany the use of chemicals to induce liver disease or viruses to mimic Hepatitis C^103,104^.

## Limitations of the study

Due to the abbreviated mortality of *Sco1* mice, we did not analyze animals past P47 which may have limited our ability to fully grasp the magnitude of their autoAb responses as well as the mechanisms leading to autoAb generation. Future studies will be performed with hepatocyte-specific *Cox10* knockout mice as these animals are more mildly affected than *Sco1* mice yet also possess increased plasma AFP, peripheral blood leukopenia and reductions in liver ATP content^24^. While the work presented here did not evaluate the immune response in the context of pathogenic infection or immunization with a model antigen, these studies will be of particular interest moving forward.

## Supporting information

Key Resources Table

## Acknowledgments

This work was supported by the Canadian Institutes of Health Research under Award Number PJ9-185660 (SCL and PDP), Saskatchewan Health Research Foundation Award Number 6230 (PDP) and the Natural Sciences and Engineering Research Council of Canada Award Number 2024-06646 (PDP). The content is solely the responsibility of the authors and does not necessarily represent the official views of any funding sources.

## Author Contributions

K.T.P, S.C.L. and P.D.P designed experiments. K.T.P., S.G., A.B., S.C.L. and P.D.P. conducted and analyzed experiments. S.C.L. and P.D.P wrote the manuscript, and all authors approved of the manuscript.

## Inclusion and Diversity

We support inclusive, diverse and equitable conduct of research.

## Supplementary Information Titles and Legends

**Table S1. Analysis of differential protein abundance in WT and Sco1 plasma using the Ray Biotech mouse L308 protein array.**

**Table S2. Metascape analysis of Gene Ontology (GO) Biological Processes enriched terms for differentially abundant proteins in WT and Sco1 plasma.**

## STAR Methods

### RESOURCE AVAILABILITY

#### Lead Contact

Further information and requests for resources and reagents should be directed to and will be fulfilled by the Lead Contact, Peter Dion Pioli.

#### Materials availability

Materials underlying this article will be shared by the lead contact upon request.

#### Data and code availability

- Flow cytometry data reported in this study will be shared by the lead contact upon request.
- Any information required for data reanalysis is available from the lead contact upon request.

## EXPERIMENTAL MODEL AND SUBJECT DETAILS

### Experimental Animals

Homozygous *Sco1* floxed (*Sco1*^Fl/Fl^) mice^25^ were crossed with animals expressing Cre recombinase driven by the *Albumin* enhancer/promoter (B6.Cg-*Speer6*-*ps1^Tg^*(Alb–Cre)*^21Mgn^*/J, The Jackson Laboratory, Strain# 003574). F1 progeny were backcrossed onto *Sco1*^Fl/Fl^ mice to obtain F2 litters containing *WT* and *Sco1* mice, with the latter genotype lacking *Sco1* expression in hepatocytes. P47 female and male mice were used for all experiments. Animal care and use were conducted according to the guidelines of the USask University Animal Care Committee Animal Research Ethics Board.

## METHOD DETAILS

### Isolation of BM, SPL,THY, Liver and Plasma

All tissues were processed and collected in calcium and magnesium-free 1x Dulbecco’s phosphate buffered saline (PBS) in 6-cm dishes. SPL and THY were dissected and crushed between the frosted ends of two slides. BM was isolated from both femurs and tibias by cutting off the end of bones and flushing the marrow using a 26-gauge needle. Following dissection, liver was cut into smaller pieces and ∼120 mg of tissue was used per sample. The plunger from a 5 mL syringe was used to crush liver pieces and cell suspensions were passed directly through a 40-μm filter and collected in a 50 mL conical tube. Cell suspensions were centrifuged for 5 minutes at 4°C and 600g. Red blood cells were lysed by suspending cells in 3 mL of 1x red blood cell lysis buffer on ice for ∼3 minutes. Lysis was stopped with the addition of 7 mL of 1x PBS. Cell suspensions were passed through 40-μm filters and counted with a hemocytometer loaded into a Contess 3 (ThermoFisher Scientific) using Trypan Blue to exclude dead cells. Cells were centrifuged as before and cell pellets were resuspended in 1x PBS + 0.1% bovine serum albumin (BSA) at a concentration of 2×10^7^ live cells/mL and cell suspensions were maintained on ice until use.

Whole blood was obtained via cardiac puncture with a 27G needle fixed to a tuberculin syringe (BD Biosciences, Cat# 305620) and immediately transferred to EDTA-lined BD Microtainer blood collection tubes (BD Biosciences, Cat# 365992) followed by inversion mixing to avoid coagulation and hemolysis. Following a 10-minute incubation on ice, blood was centrifuged at 1000g for 5 minutes at room temperature (RT). The plasma (top clear layer) was transferred to a new tube and stored at -80°C until needed.

### Immunostaining

All staining procedures were performed in 1x PBS + 0.1% BSA. All samples were labeled with a CD16/32 antibody to eliminate non-specific binding of antibodies to cells via Fc receptors. All antibodies utilized are listed in the Key Resources Table. Samples were incubated on ice for 30 minutes in the dark with the appropriate antibodies and eBioscience Fixable Viability (Live-Dead) Dye eFluor 780 (ThermoFisher Scientific, Cat# 65-0865-14) to assess the dead cell content. The stock solution was diluted 1:250 in 1x PBS and 10 μL was added to ∼5 x 10^6^ cells per stain. Unbound antibodies were washed from cells with 1x PBS + 0.1% BSA followed by centrifugation for 5 minutes at 4°C and 600g. Supernatants were decanted, and cell pellets were resuspended in an appropriate volume of 1x PBS + 0.1% BSA + 2 mM EDTA for flow cytometric analysis. Before analysis, cells were strained through a 40-μm filter mesh and kept on ice in the dark.

### Flow Cytometry

Flow cytometry was performed on a CytoFLEX (Beckman Coulter) located in the Cancer Cluster at USask. Total cells were gated using side scatter area (SSC-A) versus forward scatter (FSC-A) area. Singlets were identified using sequential gating of FSC-height (H) versus FSC-A and SSC-H versus SSC-A. All data were analyzed using FlowJo (v10) software.

### Detection of Liver AutoAbs

Plasma from P47 *WT* and *Sco1* female and male littermates was used for flow cytometry analysis of autoantibody binding of liver cells. ∼1×10^6^ liver cells (50 μL) were aliquoted per test into 5 mL tubes on ice containing 50 μL of 1X PBS + 0.1% BSA. Samples were incubated with CD16/32 antibody to block cell surface Fc receptors and Live-Dead staining reagent on ice for 30 minutes in the dark. Samples were washed with 1x PBS + 0.1% BSA, and centrifuged for 5 minutes at 600g and 4°C. Supernatant was decanted and cell pellets were resuspended in residual buffer. For intracellular autoAb staining, the eBioscience FoxP3/Transcription Factor Staining Buffer Set (ThermoFisher Scientific, Cat# 00552300) was used as follows. 500 μL of freshly prepared FoxP3 buffer 1 (1 part concentrated Fixation/Permeabilization buffer + 3 parts Fixation/Permeabilization diluent) was added per sample followed by incubation for 30 minutes in the dark at RT. 1 mL of FoxP3 buffer 2 (1 part concentrated Permeabilization buffer + 9 parts nuclease-free H_2_0) was added and samples were centrifuged for 5 minutes at 600g and 4°C. Supernatants were decanted and samples resuspended in residual buffer. All samples were stained once more with a CD16/32 antibody to block intracellular Fc receptors. Additionally, liver samples were treated with an αDNA antibody as a positive control (ThermoFisher Scientific, Cat# MA1-10600) or various amounts of *WT* and *Sco1* plasma for 30 minutes in the dark at RT. 1 mL of FoxP3 buffer 2 was added and samples were centrifuged as above. To detect bound antibodies, samples were incubated with FITC-conjugated Igκ and Igλ antibodies for 30 minutes in the dark at RT. 1 mL of FoxP3 buffer 2 was added and samples were centrifuged as above. A final wash of 2 mL of 1x PBS + 0.1% BSA was added, and samples were centrifuged as before. Supernatants were decanted, cell pellets were resuspended in an appropriate volume of 1x PBS + 0.1% BSA + 2 mM EDTA and samples were kept on ice and in the dark until flow cytometric analysis. All data are shown following background subtraction of samples stained only with Igκ+Igλ-FITC Abs.

### Pre-B Colony Assay

Pre-B cell colony forming potential of BM from P47 *WT* and *Sco1* female and male littermates was assayed by mixing 10^5^ total BM cells in 1 mL of methylcellulose (MC) medium. MC medium was prepared by supplementing phenol red-free RPMI 1640 (Gibco/ThermoFisher Scientific, Cat# 11835030) with methylcellulose (1%, MethoCult H4100, STEMCELL Technologies, Cat# 04100), heat-inactivated fetal bovine serum (30%, Gibco/ThermoFisher Scientific, Cat# 12483020), 2-Mercaptoethanol (50 μM, Thermo Scientific Chemicals, Cat# AC125472500), Penicillin-Streptomycin (100 U/mL, Gibco/ThermoFisher Scientific, Cat# 15140122), L-glutamine (2 mM, Gibco/ThermoFisher Scientific, Cat# 25030081), Gentamicin (250 μg/mL, Gibco/ThermoFisher Scientific, Cat# 15750060), sodium pyruvate (1 mM, Gibco/ThermoFisher Scientific, Cat# 11360070), MEM non-essential amino acids (1x, Gibco/ThermoFisher Scientific, Cat# 11140050), MEM vitamins (1x, Gibco/ThermoFisher Scientific, Cat# 11120052) and IL-7 (10 ng/mL, PeproTech/ThermoFisher Scientific, Cat# 2171710UG). The mixture was plated in 12-well plates. Surrounding empty wells were filled with RPMI to prevent evaporation. Plates were incubated at 37°C in a 5% CO_2_ and air incubator and colonies were counted in a blinded manner following 10 days of culture. Subsequently, 2 mL of 1x PBS was added per well and plates were incubated on an orbital shaker at 250 revolutions per minute for 30 minutes at RT. Well contents were transferred into 5 mL tubes on ice. Wells were washed with 1 mL 1x PBS and this wash was added to the previously collected cells. Cells were pelleted by centrifugation for 5 minutes at 4°C and 600g. Supernatant was discarded, and cells were resuspended in 1x PBS + 0.1% BSA. Samples were split evenly and stained for CD19 expression or left unstained to gauge background fluorescence as described above.

### ELISA – Plasma Antibodies

High binding ELISA plates (Greiner, Cat# 655081) were coated using 100 μL anti-mouse IgG/IgA/IgM (H+L) (Sigma Aldrich, Cat# SAB3701043-2MG) at 5 μg/mL in 1x PBS per well and incubated at 4°C overnight covered with plastic wrap. Coating was decanted and plates were washed 3 times with 150 μL 1x PBS + 0.1% Tween 20 (Wash Solution) per well. Subsequently, 150 μL of 1x PBS + 1% BSA + 0.1% Tween (Block Solution) was added per well and plates were blocked at RT for 2 hours. Block Solution was removed and wells washed 3 times with 150 μL of Wash Solution. Purified antibody standards (Standard Curves) and plasma samples were diluted at various concentrations in 1x PBS and 100 μL per dilution was added per well. Plates were incubated at RT for 2 hours; samples/standards were removed and wells washed 3 times with 150 μL of Wash Solution. Isotype-specific horse radish peroxidase-conjugated secondary antibodies were diluted in Block Solution and 100 μL was used per well. Secondary antibodies for IgM, IgA, IgG2b and IgG3 were diluted to a final concentration of 1:5,000 while those for IgG1 and IgG2c were diluted to 1:25,000 and 1:50,000, respectively. Following a 2-hour RT incubation, wells were washed 3 times with 150 μL of Wash Solution, then incubated for 4 minutes with 100 μL 1x TMB Substrate (ThermoFisher Scientific, Cat# 00-4201-56) per well. Enzymatic reactions were stopped with the addition of 100 μL per well of 0.16M H_2_SO_4_ (Fisher Chemical, Cat# SA431-500). Optical densities (ODs) were read at 450 nm wavelength using a BioLegend Mini ELISA Plate Reader (BioLegend, Cat# 423555). Samples and standards were analyzed following subtraction of blank wells (1x PBS) and assayed in triplicate. Plasma antibody concentrations were calculated using linear portions of standard curves and the equation of a straight line (y = mx + b) where y = average OD per sample, m = slope, x = antibody concentration and b = y-axis intercept. Only experiments with a linear standard curve R^2^ > 0.98 were considered valid.

### ELISA – Plasma ANAs

The Mouse Anti-Nuclear Antigens Total Immunoglobulins ELISA Kit (Alpha Diagnostic International, Cat# 5210) was used to analyze ANAs present in plasma from P47 *WT* and *Sco1* female and male mice. The procedure was followed according to the provided instruction manual using plasma diluted 1:100 in duplicate for each sample. Plates were covered with aluminum foil during incubation periods. ODs were read at 450 nm wavelength as above with samples and standards analyzed following subtraction of blank wells (sample diluent). Calibration curves with R^2^ > 0.98 were used to calculate ANA concentrations per plasma sample similar to *ELISA – Plasma Antibodies*. Data are presented as absorbance due to the low level of detection.

### ELISA – Plasma AFP

The Mouse AFP Quantikine ELISA Kit (R&D Systems, Cat# MAFP00) was used to quantify AFP levels in duplicate according to the manufacturer’s specifications after diluting P47 *WT* and *Sco1* plasma 1:2.5 and 1:666,700, respectively. Plates were covered with aluminum foil during incubation periods. ODs were read at 450 nm wavelength using a SpectraMax 384 plus Microplate reader (Molecular Devices), and samples and standards analyzed following subtraction of an internal OD read at 540 nm wavelength. Data for standards were further corrected by subtracting blank wells containing sample diluent, log transformed and the resultant linear regression curve with R^2^ = 0.91 was used to calculate AFP concentrations per plasma sample.

### Plasma Protein Array

P47 *WT* and *Sco1* female and male littermate plasma was collected, immediately snap frozen and then frozen at -80°C. *WT* and *Sco1* plasma was thawed, and plasma pools were generated by combining equal volumes of plasma from 4 individual mice from 2 independent pairs of male and female littermates. Plasma pools were subsequently dialyzed against 1x PBS using Slide-A-Lyzer™ MINI Dialysis Devices (3.5K MWCO, ThermoFisher Scientific, Cat# 69550) at 4°C with continuous mixing. After 3 hours, the 1x PBS was replaced, and samples were left overnight for a second round of dialysis. Dialyzed *WT* and *Sco1* plasma (38 µL) was prepared for and incubated with the Mouse L1308 Array Glass Slide kit (RayBiotech, Cat# AAM-BLG-1-4), according to the manufacturer’s instructions. The fully processed glass slide was imaged using the LI-COR Odyssey system (Software Version: 2.1.12) and processed using LI-COR Image Studio.

The intensity of the array spots was calculated based on the manufacturer’s recommendations. Briefly, each antibody target was spotted on the array in duplicate, and the mean value between duplicates was used for all subsequent calculations. The intensity of a given signal from the *Sco1* array was normalized to that of the corresponding *WT* array signal by multiplying against the ratio in the geometric means of the internal, positive controls between both arrays. The mean value from all negative control spots was then subtracted from the normalized values to yield the final signal intensity. Proteins were considered significantly altered based upon the following criteria: 1) the protein was regulated in the same direction for all 3 experiments (i.e., Up in *Sco1* or Down in *Sco1*), 2) the average non-transformed fold change (*Sco1*/*WT*) was >2.0 or <0.5 and 3) the standard deviation was <3 for the fold change from all 3 experiments.

## QUANTIFICATION AND STATISTICAL ANALYSIS

The numbers of mice used (n =) or replicates performed per experiment are listed in the Figure Legends. Quantification of cell numbers and various flow cytometry data are graphically represented as mean ± SEM. A Student’s t-Test was used for statistical comparisons between 2 groups. Statistically significant p-values are shown within each figure.

