## Supplementary material for "A hepatocyte-specific cytochrome *c* oxidase deficiency in mice leads to a lymphopenia owing to deficiencies in bone marrow progenitors": Figures S1-S5

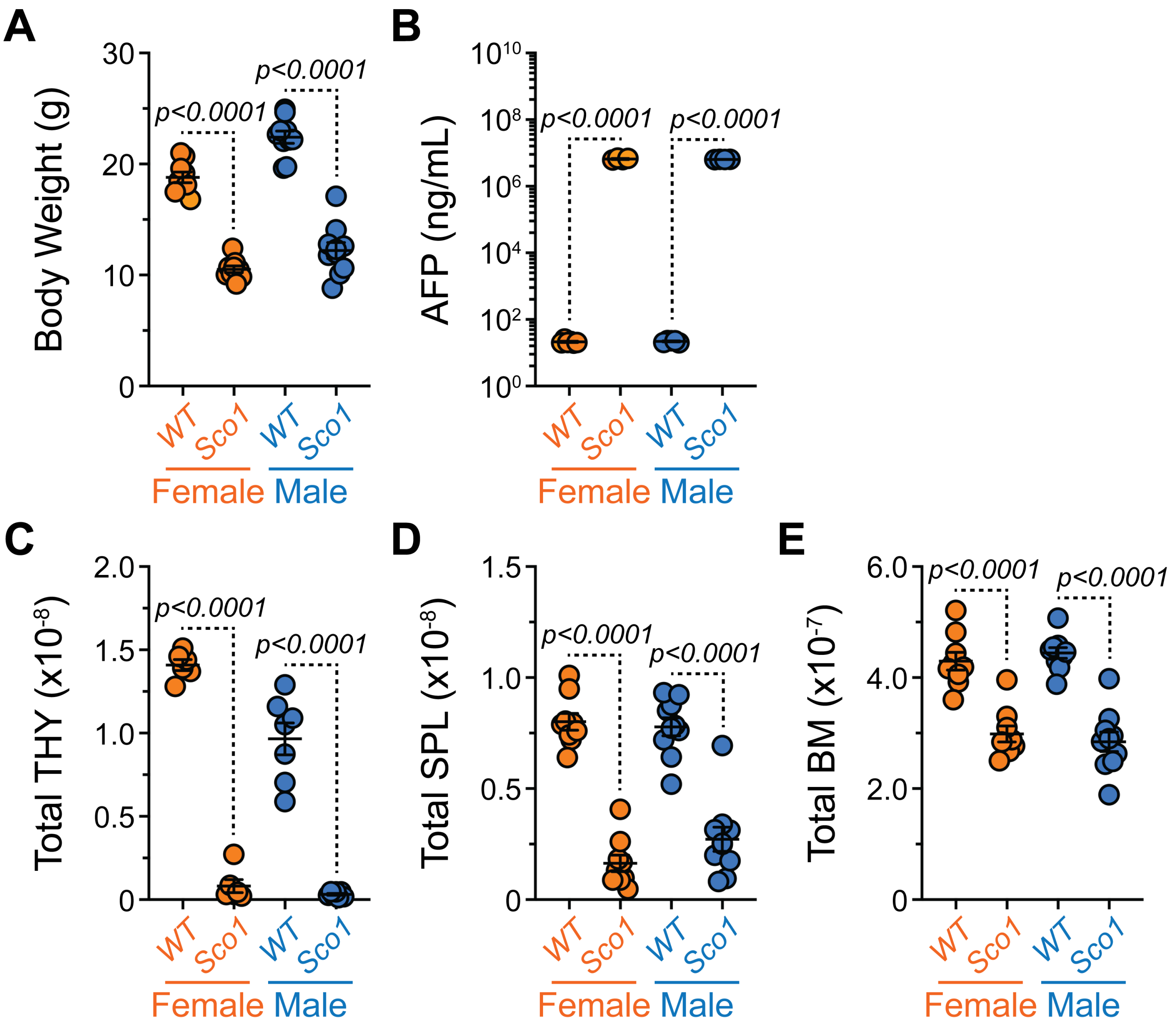


**Figure S1. P47 *Sco1* mice demonstrate reductions in lymphoid organ cellularity. Related to Figures 1-6.** (A) Body weights of *WT* and *Sco1* mice. (B) AFP levels in plasma from *WT* and *Sco1* mice. (C-E) Total cellularity of (C) THY, (D) SPL and (E) BM isolated from *WT* and *Sco1* mice. (A-E) Symbols represent individual mice. Horizontal lines represent mean ± SEM. Statistics: Unpaired Student's t-Test. (A, D-E ) *WT* Female: n = 9, *Sco1* Female: n = 9, *WT* Male: n = 10, *Sco1* Male: n = 10. (B) *WT* Female: n = 6, *Sco1* Female: n = 4, *WT* Male: n = 5, *Sco1* Male: n = 4. (C) *WT* Female: n = 6, *Sco1* Female: n = 6, *WT* Male: n = 7, *Sco1* Male: n = 7.

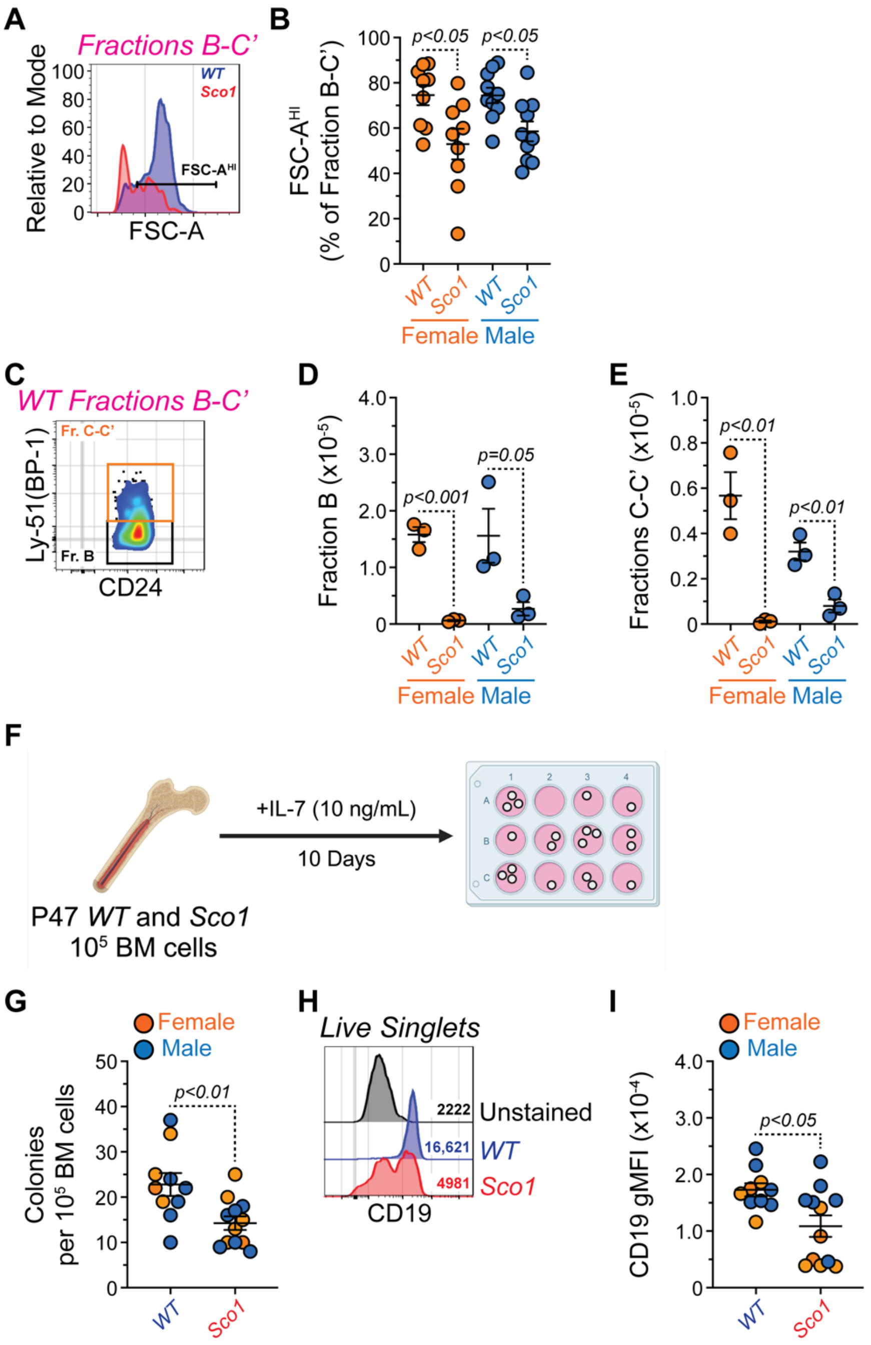

**Figure S2. BM B cell development is severely depleted in P47 *Sco1* mice. Related to Figure 4.** (A) Flow cytometry histogram overlay depicting representative gating of FSC-A$^{HI}$ cells in Hardy Fractions B-C' from *WT* and *Sco1* BM. Cells pre-gated on Hardy Fractions B-C'. (B) Percentages of FSC$^{HI}$ cells within Hardy Fractions B-C' from *WT* and *Sco1* mice. (C) Flow cytometry plots depicting representative gating of Hardy Fractions B and C-C' from *WT* BM. Cells pre-gated on Hardy Fractions B-C'. (D-E) Numbers of (D) Hardy Fraction B and (E) Hardy Fractions C-C' B cells from *WT* and *Sco1* BM. (F) Schematic depicting B cell progenitor colony assay. $10^5$ BM cells from *WT* and *Sco1* mice were plated in methyl cellulose containing IL-7 (10 ng/mL). Figure made with BioRender. (G) Numbers of colonies generated by *WT* and *Sco1* BM. (H) Flow cytometry histogram overlay depicting CD19 staining of colony assays initiated with *WT* and *Sco1* BM. Unstained cells are shown for comparison. Numbers in plot indicated CD19 gMFIs. Cells pre-gated on total live singlets. (I) CD19 surface staining gMFIs of cells from colony assays initiated with *WT* and *Sco1* BM. (B, D-E) Symbols represent individual mice. Horizontal lines represent mean ± SEM. Statistics: Unpaired Student's t-Test. (B) *WT* Female: n = 9, *Sco1* Female: n = 9, *WT* Male: n = 10, *Sco1* Male: n = 10. (D-E) *WT* Female: n = 3, *Sco1* Female: n = 3, *WT* Male: n = 3, *Sco1* Male: n = 3. (G, I) Symbols represent individual colony assay wells initiated from 3 female and male *WT* and *Sco1* mice. Horizontal lines represent mean ± SEM. Statistics: Unpaired Student's t-Test.

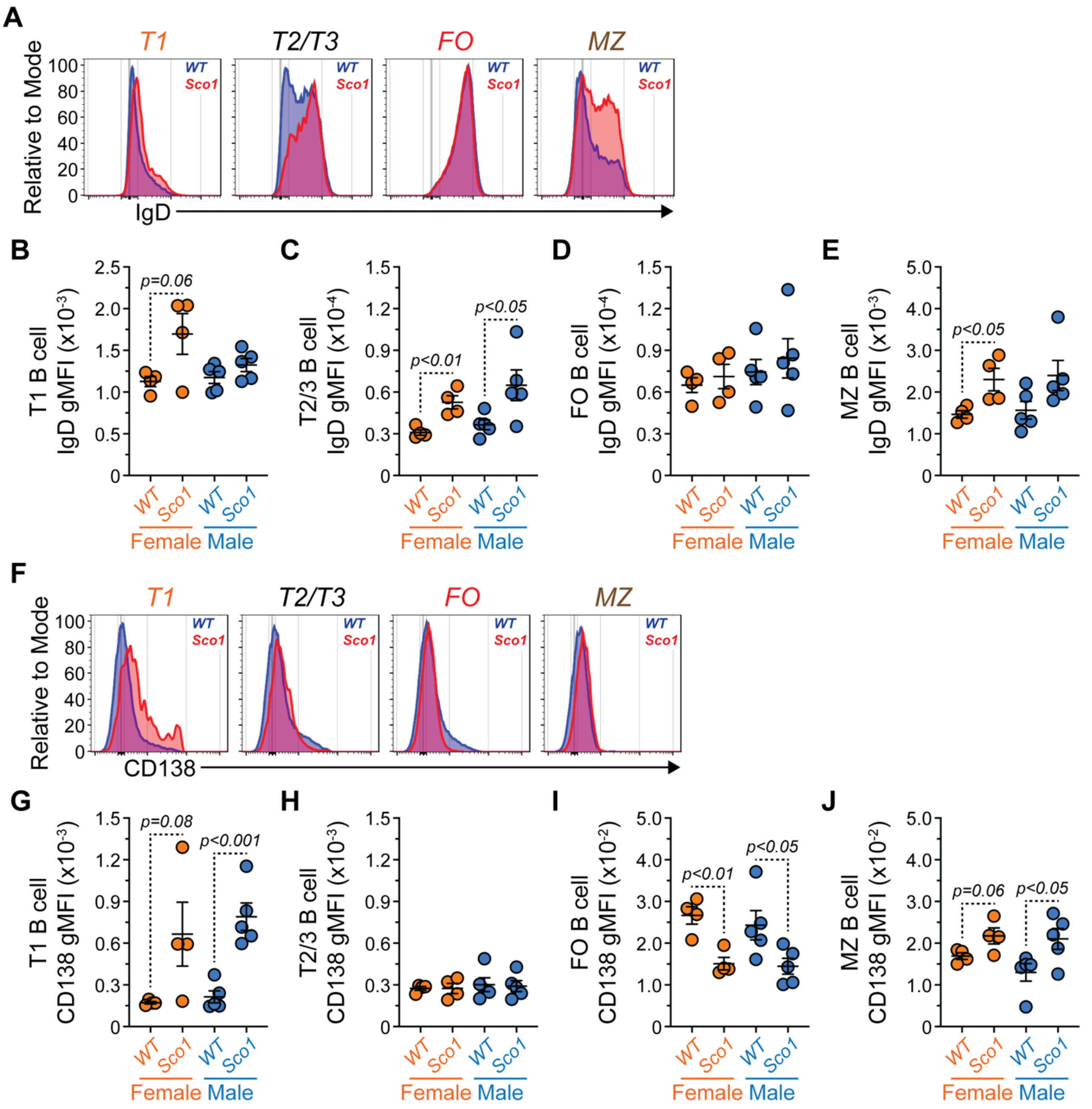


**Figure S3. SPL B cell maturation is dysfunctional in P47 *Sco1* mice. Related to Figure 5.**
(A) Flow cytometry overlays showing IgD staining on the surface of T1, T2/3, FO and MZ B cells from *WT* and *Sco1* SPL. (B-E) IgD surface staining gMFI for (B) T1, (C) T2/3, (D) FO and (E) MZ B cells from *WT* and *Sco1* SPL. (F) Flow cytometry overlays showing CD138 staining on the surface of T1, T2/3, FO and MZ B cells from *WT* and *Sco1* SPL. (G-J) CD138 surface staining gMFI for (G) T1, (H) T2/3, (I) FO and (J) MZ B cells from *WT* and *Sco1* SPL. (B-E, G-J) Symbols represent individual mice. Horizontal lines represent mean ± SEM. *WT* Female: n = 4, *Sco1* Female: n = 4, *WT* Male: n = 5, *Sco1* Male: n = 5. Statistics: Unpaired Student's t-Test.

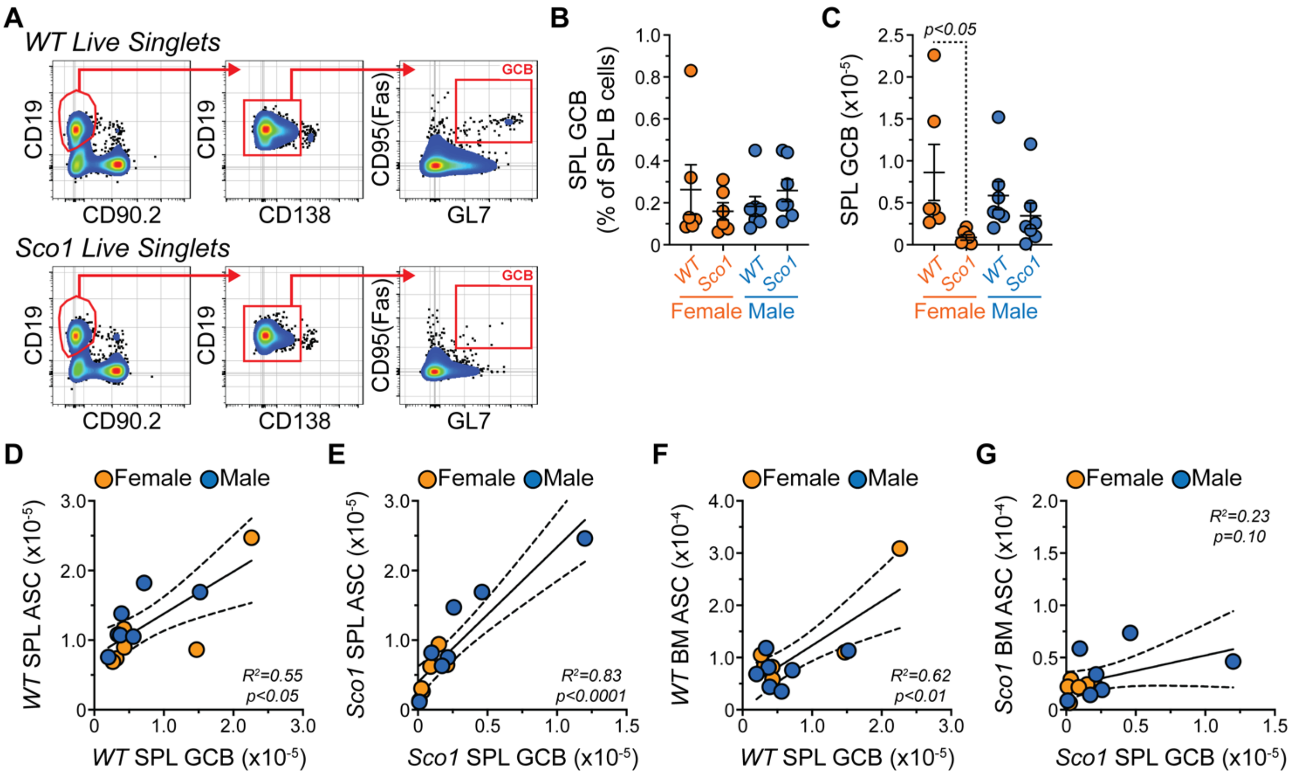


**Figure S4. GCBs are reduced in the SPL from P47 *Sco1* mice. Related to Figure 6.**
(A) Flow cytometry plots depicting representative gating of GCBs in the *WT* and *Sco1* SPL. Cells pre-gated on total live singlets. (B) Percentages of GCBs within B cells from *WT* and *Sco1* SPL. (C) Numbers of GCBs from *WT* and *Sco1* SPL. (D-G) Correlation of (D) *WT* SPL ASCs and SPL GCBs, (E) *Sco1* SPL ASCs and SPL GCBs, (F) *WT* BM ASCs and SPL GCBs and (G) *Sco1* BM ASCs and SPL GCBs. (B-C) Symbols represent individual mice. Horizontal lines represent mean ± SEM. *WT* Female: n = 6, *Sco1* Female: n = 6, *WT* Male: n = 7, *Sco1* Male: n =7. Statistics: Unpaired Student's t-Test. (D-G) Symbols represent individual mice. Lines indicate means and 95% confidence intervals. *WT* Female: n = 6, *Sco1* Female: n = 6, *WT* Male: n = 7, *Sco1* Male: n =7. Statistics: Linear regression analysis.

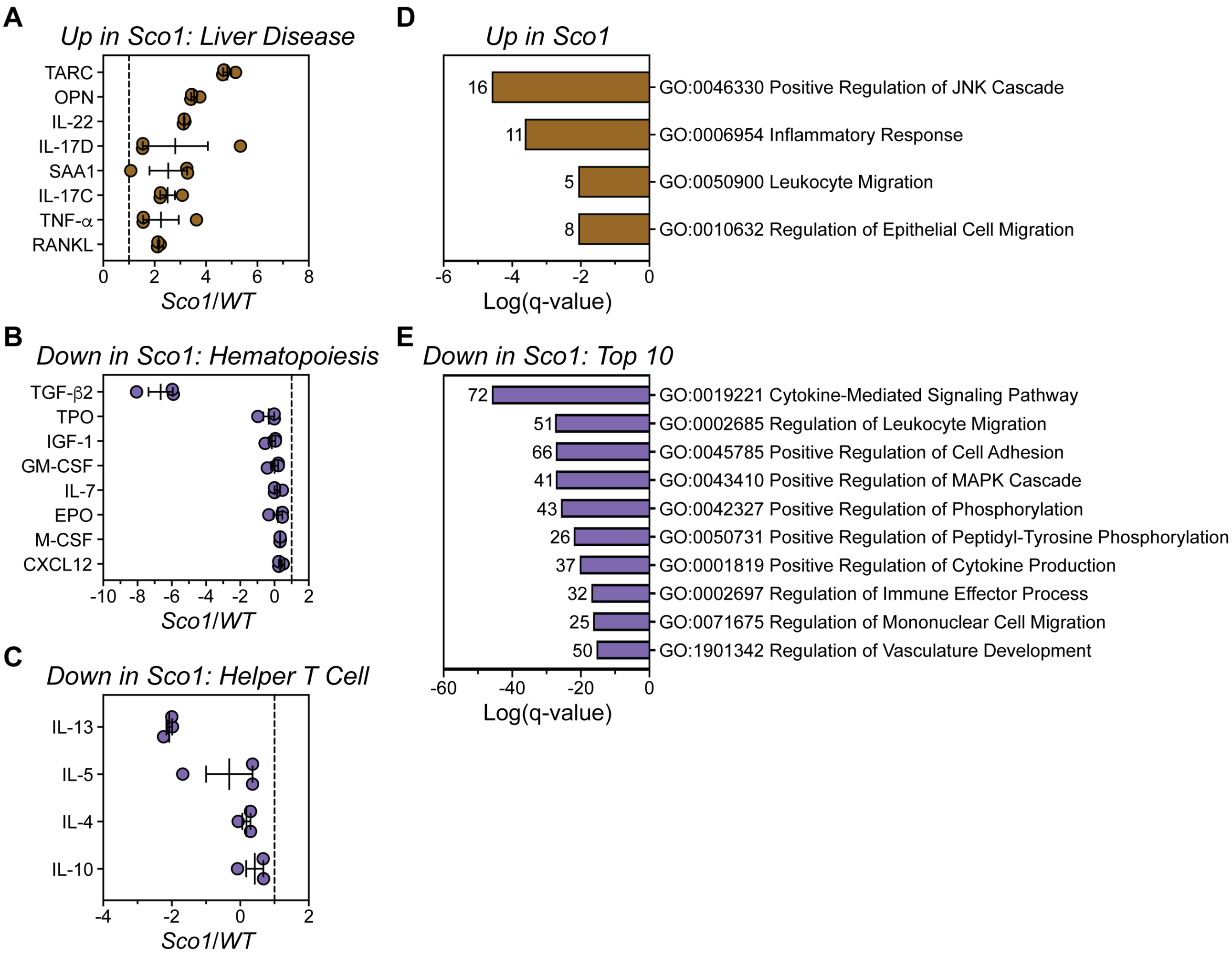


**Figure S5. Protein abundance in P47 *Sco1* plasma is altered compared to *WT* plasma. Related to Tables S1-S2.**

(A) Relative abundance (*Sco1*/*WT*) of selected proteins that are increased in *Sco1* plasma and associated with liver disease. (B,C) Relative abundance (*Sco1*/*WT*) of selected proteins that are decreased in *Sco1* plasma and associated with (B) hematopoiesis or (C) helper T cell functions. (A-C) Dashed line indicates no change in expression. (D-E) Log(q-value) statistical significance for GO categories associated with (D) proteins increased in *Sco1* plasma and (E) proteins decreased in *Sco1* plasma. Only the top 10 GO categories are shown for (E). Numbers in plots indicate the number of proteins associated with each GO. Values derived from Metascape analyses presented in Table S2.
